# Hydration-induced structural transitions in biomimetic tandem repeat proteins

**DOI:** 10.1101/2021.01.12.426322

**Authors:** Romeo C. A. Dubini, Huihun Jung, Melik C. Demirel, Petra Rovó

**Affiliations:** Department of Chemistry and Pharmacy, Ludwig-Maximilians-Universität München, 81377 Munich, Germany; Center for Nanoscience (CeNS), Faculty of Physics, Ludwig-Maximilians-Universität München, Schellingstraβe 4, 5th floor, 80799 Munich, Germany; Center for Research on Advanced Fiber Technologies (CRAFT), Materials Research Institute, Department of Engineering Science and Mechanics, and Huck Institutes of Life Sciences, Pennsylvania State University, University Park, Pennsylvania 16802, United States of America

## Abstract

A major challenge in developing biomimetic, high-performance, and sustainable products is the accurate replication of the biological materials’ striking properties, such as high strength, self-repair, and stimuli-responsiveness. The rationalization of such features on the microscopic scale, together with the rational design of synthetic materials, is currently hindered by our limited understanding of the sequence-structure-property relationship. Here, employing state-of-the-art nuclear magnetic resonance (NMR) spectroscopy, we link the atomistic structural and dynamic properties of an artificial bioinspired tandem repeat protein TR(1,11) to its stunning macroscopic properties including high elasticity, self-healing capabilities, and record-holding proton conductivity amongst biological materials. We show that the hydration-induced structural rearrangement of the amorphous Gly-rich soft segment and the ordered Ala-rich hard segment is the key to the material’s outstanding physical properties. We found that in the hydrated state both the Ala-rich ordered and Gly-rich disordered parts contribute to the formation of the nanoconfined *β*-sheets, thereby enhancing the strength and toughness of the material. This restructuring is accompanied by fast proline ring puckering and backbone *cis-trans* isomerization at the water-protein interface, which in turn enhances the elasticity and the thermal conductivity of the hydrated films. Our in-depth characterization provides a solid ground for the development of next-generation materials with improved properties.

## Introduction

Self-assembling proteins are valuable building blocks allowing the manufacturing of materials with versatile physicochemical properties and functions based on their secondary and tertiary structures. Well-studied motifs from tandem repeat proteins—such as silk, elastin, collagen, keratin, resilin, and squid ring teeth (SRT)—have been frequently used to develop multifunctional materials for diverse applications.^1–5^ Transcriptome analysis of SRT proteins revealed the presence of several segmented copolymer protein sequences with alternating ordered and amorphous regions and overall molecular weight ranging from 15 kDa to 55 kDa.^6^ Compared to other biopolymers (e.g., keratin, resilin, and elastin), the proteinaceous structure of the SRT does not have any covalent cross-linker and dissolves in organic solvents such as hexafluoro-2-propanol (HFIP) or dimethyl sulfoxide (DMSO) (with solubility greater than 100 g/L) or weak acidic/alkaline buffers (albeit with lower solubility, ~1 g/L), while it reversibly aggregates whenever the solvent is removed. The protein’s amorphous regions include flexible chains rich in glycine, leucine and tyrosine, while the ordered regions are formed by Ala-rich segments stabilized by tight packing of alternating side chains, separated by proline residues. Based on consensus sequences derived by inspection of the native SRT proteins of squid species Loligo vulgaris, we recently constructed a series of DNA sequences encoding SRT-inspired polypeptides.^7,8^ A representative amino acid sequence consisted of an ordered segment of PAAASVSTVHHP, a disordered segment of YGYGGLYGGLYGGLGYG, as well as a cleavage side of STGTLS that aided the gene-assembly of the tandem repeat proteins (Fig. 1a). These recombinant tandem repeat biopolymers—abbreviated as TR(*f, n*)—can be tuned for predefined physical properties by controlling their network morphology by varying the ratio of the ordered and disordered segments (*f*) or the overall length of protein by increasing the repeat numbers (*n*).^9^ The biopolymer can be designed to exhibit a variety of unusual, stimuli-responsive physical features including rapid and reversible hydration-dependent switching in thermal conductivity,^10^ highest proton conductivity among proteinaceous materials^11^ and exceptionally fast self-healing properties upon heating.^12,13^ A detailed structural analysis is essential to understand the atomistic origin of these unique features. In addition, insights into the nanoscale architecture and hierarchical organization could support the development and fine tuning of future bio-inspired green materials.^7,9,14–16^ However, the most commonly applied spectroscopic techniques—e.g. Fourier-transform infrared (FTIR), Raman, and circular dichroism spectroscopy—provide only coarse-grained information about the conformational distribution of these materials. Structural proteins are inherently heterogeneous, presenting a combination of *α*-helix, *β*-sheet and *β*-turn arrangements embedded into an amorphous, glycine-rich random-coil matrix, making their characterization challenging via X-ray diffraction or electron microscopy techniques. One of the methodologies that has the potential to provide atomic resolution structural details about the material’s assembly is nuclear magnetic resonance (NMR) spectroscopy. Here, the conformation-dependent chemical shifts allow for sensitive monitoring of the change in local structure and electronic environment (H-bonding, *π–π* stacking, lattice packing) on an amino acid-specific manner. Moreover, it provides information about potential interaction partners and local dynamics occurring from the picoseconds up to the seconds time scale. NMR has thus been widely and successfully employed to elucidate the structure and dynamics of natural fibers such as collagen, elastin, silks from spiders, silkworms and other insects, and that of man-made, bioinspired synthetic analogues.^17–19^ In this study, we present the atomic resolution structural characterization of recombinantly produced SRT-inspired tandem repeat protein TR(1,11), i.e. for a repeat protein where the ratio between the amorphous and ordered segment is approximately one, and this 35 residues long sequence is repeated 11 times (see Supplementary Methods for the exact sequence). The recombinant protein is straightforward to produce in isotopically enriched forms, enabling multidimensional heteronuclear NMR studies. We use solution-state NMR to gain information about TR(1,11) in a random coil, fast-tumbling state, where the well-resolved signals enable residue-specific resonance assignments which in turn give insights into the conformational preference of the backbone dihedral angles. In addition, we investigate the overall and local rotational dynamics of the dissolved protein. Besides, we employ fast magic angle spinning (MAS) solid-state NMR to assess the structural assembly and the hydration-induced conformational and motional changes occurring to TR(1,11). In this endeavor, we consider three solid-state variants: the ambient powder (AP), ambient film (AF), and hydrated film (HF) forms (Fig. 1b). The comprehensive analysis and comparison of these conditions enable the rationalization of the exceptional physical properties of TR biopolymers.

**Figure 1.**
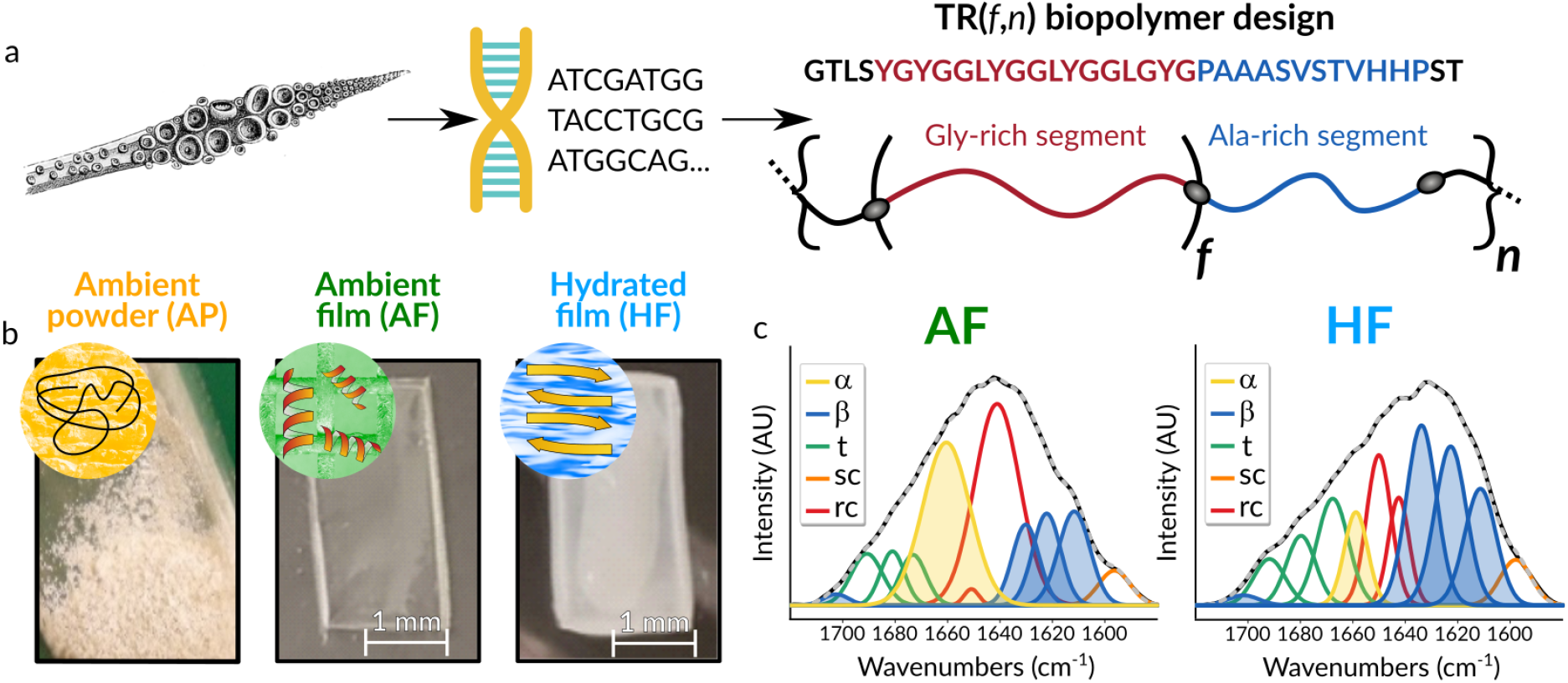
(a) Transcriptome analysis of squids’ ring teeth proteins led to the design of the artificial Tandem Repeat TR(*f, n*) proteins.^5,10,13^ (b) Introduction of the three solid-state samples considered: ambient powder (AP), ambient film (AF) and hydrated film (HF). (c) Deconvoluted FTIR spectra for samples AF and HF. Each band is labelled as *β*-sheet (*β*), *α*-helix (*α*), turn (t), side chain (sc) or random coil (rc) according to the spectral regions of amide I (1600–1700 cm^-1^). Shaded areas under α and β curves highlight the large structural rearrangement upon SRT film hydration, i.e. AF → HF transition.

## Results

### Protein design and coarse-grained assessment

Fig. 1a shows the general strategy of SRT-inspired tandem repeat protein (TR) design. We prepared the gene constructs and vectors of TR(1,11), and then expressed and purified the encoded proteins in recombinant *E. coli* in a pET-derived system according to our previously published methods.^8^ To confirm that the produced protein has the expected molecular weight of 39.2 kDa, we performed matrix-assisted laser desorption/ionization time-of-flight spectrometry (Fig. S1). We kept TR(1,11) either in a freeze-dried powder form or dissolved it in HFIP and cast as a thin film (Fig. 1b). When we washed the protein films with deionized water, a structural transition became apparent from the change in the film’s transparency (i.e., freshly cast films are clear, whereas hydrated films are opaque, Fig. 1b). To assess the coarsegrained structural transitions upon film hydration, we first employed FTIR spectroscopy (Fig. 1c and Fig. S2), and analyzed the amide I bands by Fourier self-deconvolution (FSD) and Gaussian fitting. Following the literature of fibrous proteins, we assigned the FTIR bands to secondary structures, and used the integrals to estimate the relative fraction of the various secondary structure elements present in the ambient and hydrated films.^20^ On a qualitative level, this analysis revealed a substantial increase in *β*-sheet content alongside with the almost complete disappearance of α-helical structures (Fig. 1c and Table S1).

### Solution-state NMR studies of TR(1,11)

Overall, two sets of assignment spectra were collected, the former immediately after powder dissolution and 3-hours sonication, and the latter after ~60 days of incubation at room temperature. To assess whether individual segments of the protein displayed any preference for *α*-helical or *β*-sheet conformations, we analyzed their chemical shifts in the freshly prepared and later in the matured samples. In addition, in-between the two assignment data sets, we performed ^15^N relaxation experiments reporting on the ps-ns time-scale dynamics in order to gain a quantitative understanding of the protein motions in disordering conditions. A first inspection of the peaks in the ^1^H–^15^N HSQC spectrum (Fig. 2a) suggests that the protein tumbles rapidly in solution and behaves as an intrinsically disordered protein. The disordered nature of the sample is apparent from the relatively narrow distribution and narrow line-widths of the amide ^1^H and ^15^N resonances, and it was later corroborated by the quick tumbling time derived from the relaxation data (*vide infra*). The sequential degeneracy of TR(1,11) results in a considerable simplification of the NMR spectra since the 11 subunits, each containing 35 amino acids, lead to degenerate chemical shifts for each equivalent residue, suggesting an identical chemical environment and thus indistinguishable structural propensities for each repeat unit. The assignment of the ^1^H, ^15^N, and ^13^C resonances followed standard procedures using triple resonance experiments. For residues A23, A25, and S26 we identified multiple signals belonging to the same residues. Inspection of the hCCcoNH stripes for A23 and A23’ (Fig. 2b) established that such feature arises from the slow *cis-trans* isomerization of the G21–P22 peptide bond. Reassuringly, the intensity of the second sets of peaks was found to be consistent with a ratio of ~30:70 (*cis:trans*), in agreement with the expected Boltzmann equilibrium distribution of a freely rotating Gly–Pro peptide bond.^21,22^ To study the structural preference of the solubilized TR(1,11), we evaluated its ^13^C chemical shifts. The difference between the Δ*δ*C*α* and Δ*δ*C*β*, where Δ*δ*C*i* = *δ*C_obs_ – *δ*C_rc_ is the deviation between the observed “obs”) and the random coil (“rc”) chemical shifts of the *i*^th^carbon, is an accurate indicator of transient *α*-helix or *β*-strand formations. In case a consecutive stretch of more than three amino acids shows the same trend above a certain threshold (e.g. ±2 ppm), then that segment can be associated with *α*-helical (+2 ppm) or *β*-strand (−2 ppm) preference.^23^ According to this analysis, the PAAA segment possesses a clear propensity towards *β*-strand conformation apparent from the < –2 shift differences for the corresponding ^13^C chemical shifts (Fig. 2c). No other amino acid stretches satisfied the above criteria, albeit they all displayed a moderate tendency to *β*-strands formation. A second aspect emerging from the analysis of ^13^C resonances is the large chemical shift perturbation experienced by H31 and H32 in the time elapsing the two experimental sets (Fig. 2c). Such alteration is mirrored in the ^1^H–^15^N correlation spectrum, where resonances V30, H31, and H32 witness a chemical shift perturbation in the order of ~0.5 ppm (Fig. 2d). The ^13^C secondary chemical shift changes suggest that the initial extended backbone conformation of H31, H32 over time transformed more towards random coil.

**Figure 2.**
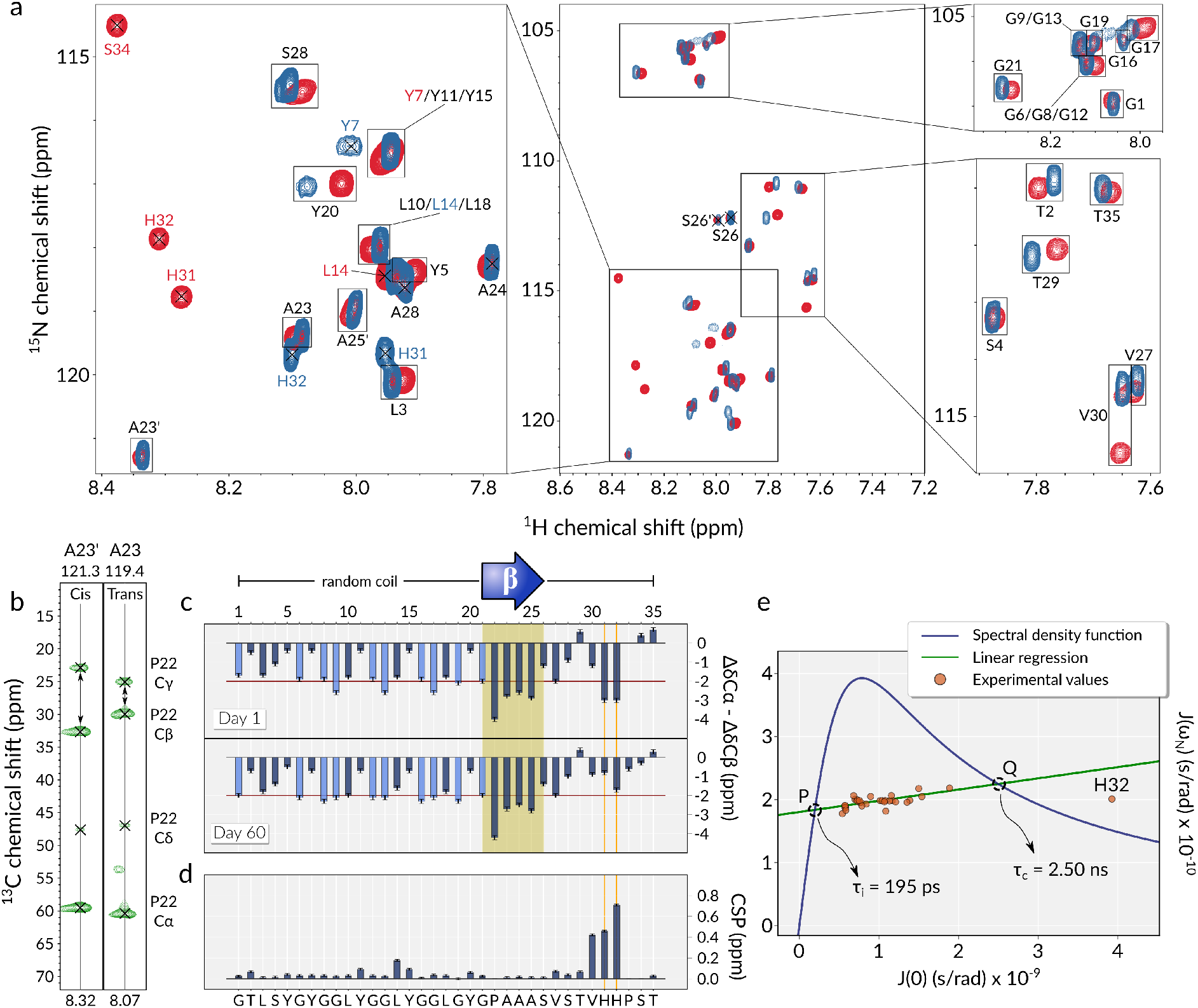
(a) ^1^H–^15^N HSQC spectra of ^15^N labeled TR(1,11) after dissolution in DMSO-*d*_6_ and sonication (blue) and after ~60 days of incubation at room temperature (red) measured at 800 MHz ^1^H Larmor frequency. Black assignment labels refer to both spectra. (b) hCCcoNH A23 and A23’ strip plots providing chemical shift of the *cis* (left) and *trans* (right) isomers for residue P22. (c) Δ*δ*C*α* – Δ*δ*C*β* chemical shifts reflect structuring propensities in TR(1,11). Gly Δ*δ*C*α* values are displayed in light blue. Yellow shade indicates a segment with *β*-sheet structural preference. Vertical orange lines mark the sites which change conformation during the 60 days of incubation. (d) ^1^H–^15^N chemical shift perturbation of TR(1,11) up on 60 days incubation at room temperature. (e) Correlation plot of the *J*(0) and *J*(*ω*_N_) spectral density values derived from the ^15^N relaxation rates. Solid green line marks the linear fit of the obtained values. Solid blue line indicates the limiting solution with a single correlation time with a Lorentzian form for the spectral density function. The intercepts *P* and *Q* correspond to the correlation time for internal (*τ*_i_) and global (*τ*_c_) tumbling times, respectively. The errors of the derived J values are smaller than the symbol size.

### Diffusion and dynamics measurements

To test whether the observed time-dependent structural change in the VHHP region is the consequence of aggregation, we compared ^1^H diffusion ordered spectroscopy (DOSY) spectra measured before and after the incubation period. The two spectra yielded nearly identical results, proving that the macroscopic diffusional behavior of TR(1,11) is not time-dependent in the dilute regime, and hence the observed structural change in the VHHP region is not the result of coagulation. Based on the experimentally determined diffusion coefficient, we estimated the hydrodynamic radius of the solubilized TR(1,11) to be 33.0 Å (see SI Text for more details), a value that fits well within the theoretical limits for a protein of equal size in the completely folded and unfolded conformations (*R_h_* = 27.0 Å and *R_h_* = 67.2 Å, respectively).^24^ Finally, in order to probe the backbone dynamics of TR(1,11) in solution state, we measured ^15^N *R*_1_, *R*_2_ rates, and ^1^H–^15^N heteronuclear nOe ratios and analyzed them within the framework of the reduced spectral density mapping (RSDM) formalism, allowing for a complete, model-independent assessment of the dynamics of the system.^25^ The analysis revealed a linear correlation between the experimentally obtained *J*(0) and *J*(*ω*_N_) spectral density values (Fig. 2e, green line), implying that the motion of each amide site can be accurately described by an overall tumbling time of 2.5 ns, and with an internal bond fluctuation that occurs with a correlation time of 195 ps. A folded protein of similar molecular weight (~40 kDa) is expected to tumble an order of magnitude slower.^26^ The only outlier identified in the correlation plot was H32, whose elevated *J*(0) indicates substantial *μ*s-ms time-scale exchange contribution to the R_2_ rates (Fig. S3).

### Solid-state NMR studies of AP, AF, and HF

After scrutinizing the solution-state behavior of TR(1,11), we turned to its solid forms and monitored the structural and dynamical changes induced by film casting and subsequent hydration using fast MAS solid-state NMR spectroscopy with spinning frequency of 55.55 kHz. We recorded 1D and 2D spectra of the uniformly ^13^C, ^15^N labeled AP, AF, and HF TR(1,11) samples. In particular, we assessed the structural variations by following the resonance shifts in ^13^C-detected ^13^C–^13^C as well as in ^1^H-detected ^1^H–^13^C 2D correlation experiments using both dipolar-coupling-based cross-polarization (CP) and scalar-coupling-based INEPT as a mean for heteronuclear magnetization transfer, and dipolar recoupling enhanced by amplitude modulation (DREAM) or radiofrequency driven dipolar recoupling (RFDR) for homonuclear mixing. In the CP-based experiments only those signals appear which belong to the relatively rigid segments of the protein which do not exhibit large-amplitude, fast ps-ns time-scale motion that would inhibit the dipolar-coupling-based polarization transfer. Conversely, in the INEPT-based experiments only such sites lead to observable signals that have long coherence lifetimes which occurs primarily to highly mobile parts of the molecule, such as methyl side-chains or fully dissolved segments. In the unfortunate case of large amplitude intermediate (*μ*s-ms) time-scale conformational exchange, the signals would become undetectable in both CP and INEPT-based experiments due to severe exchange broadening. In such scenario, measurements at distinctly different temperatures could help observing the exchange-broadened resonances. Fig. 3 displays the 1D ^1^H, ^13^C, and ^15^N spectra of the AP, AF, and HF samples. The broad line widths indicate large structural heterogeneity which improves as TR(1,11) is cast into a dry film and further when it is hydrated. Hydration-induced line narrowing is a known phenomenon for soft biopolymer materials.^27,28^ The sharp signals that appear in the ^1^H 1D spectrum of HF belong to water as well as to added detergent and co-solvent resonances (Triton-X and HFIP). The extremely broad line widths of the ^15^N resonances inhibited their residue-specific analysis; only the Gly amide ^15^N and His side-chain ^15^N*δ* signals could be unambiguously differentiated from the rest of the bulk amide ^15^N signals. Increasing the dimension of the ^15^N spectra by recording them in a ^1^H-detected 2D fashion did not improve the ^15^N resolution (Fig. S4), hence we refrained from any further analysis of the amide resonances and focused on the carbon sites. The direct excitation ^13^C 1D spectra recorded with a recycle delay of 25 s revealed substantial changes in relative resonance positions and peak intensities among the studied samples, with largest variations at the aliphatic region (0 – 60 ppm). To increase the resolution we recorded 2D versions of the ^13^C spectra both as homonuclear ^13^C–^13^C and heteronuclear ^1^H–^13^C experiments. Besides, we recorded solution-state-like ^1^H–^13^C HSQC experiments, as well as 2D ^13^C–^13^C spectra where the initial magnetization transfer was a refocused INEPT step, and the mixing was achieved via RFDR using the pulse sequence displayed in Fig. S5. Fig. 4 and S6 compares the aliphatic and carbonyl regions of the ^13^C–^13^C DREAM spectra of the three samples collected with 4 ms mixing period and a carrier offset at 52.0 ppm. The tentative assignment was made possible by comparing the observed ^13^C chemical shifts to random coil values from the literature^29^ (Table 1), and by comparing the solution-state and solidstate ^13^C resonances. A synthetic ^13^C–^13^C solution-state spectrum, color coded by the amino acid types, is overlaid on the DREAM spectrum of the powder sample in Fig. 4a. To the most informative chemical shifts belong the Ala C*α*, C*β*, and CO and Gly C*α* and CO carbons which can be unambiguously assigned based on their unique position in the ^13^C–^13^C correlation spectra. Moreover, these chemical shifts are highly correlated with the secondary structure in which the corresponding amino acids are located and thus they could infer to the structure of the crystalline and amorphous domain of the repeat protein. As expected, the freeze-dried sample displays extremely broad ^13^C resonances (~500 Hz line width) indicating a heterogeneous sample without any local or long-range order (Fig. 4a). The Ala C*α*/C*β* crosspeaks appear at all possible resonances suggesting a wide distribution of backbone torsion angles (Fig. 5, orange spectrum); Gly and most other amino acids are in random coil conformation. When the sample is cast into a film, the local order increases which is apparent from the peak sharpening. In the AF form, Ala resonances appear exclusively in an *α*-helical conformation (C*α*/C*β*: 51.7/14.0 ppm, Fig. 5, green spectrum) while Gly has random coil or extended conformations (C*α*/CO: 43.4/171.6 ppm, Fig. 4b). In agreement with the observations of the FTIR study, we observe that hydration induces large structural rearrangements, with the most striking change being the almost complete transformation of the Ala conformation from *α*-helical to extended *β*-strand apparent from the shift of the C*α*/C*β* resonances from 51.7/14.0 ppm to 47.8/19.5 ppm, only ~10 % of the alanines remain in helical form (Fig. 5 cyan spectrum). Meanwhile, Gly C*α* shifts upfield, from 43.4 to 41.6 ppm, and the CO resonance splits into two populations, one at 170.3 ppm and another one at 168.2 ppm with an approximate ratio of 40%: 60%; based on their relative chemical shifts, both of these populations represent extended structures but apparently in different electronic environments (Fig. S7). The extreme upfield shifted Gly C*α* resonance with respect to the random coil value could indicate 3_1_-helical conformation in the amorphous region, however the Gly CO shift does not match the corresponding shifts observed in 31-helical (AGG)10 model peptides.^30^ Besides Gly and Ala, a few other amino acids also displayed substantial, hydration-dependent shifts in their ^13^C resonances (Table 1 and S2). These residues include Pro, Ser, and Thr, all of which transitioned from predominantly *α*-helical to extended conformations as apparent from the shift of Pro C*α*/C*β* cross-peak from 63.4/28.7 ppm to 61.3/29.3 ppm, Ser C*α*/C*β* cross-peak from 59.0/59.3 ppm to 53.7/62.8, and Thr C*α*/C*β* cross-peak from 63.2/66.2 ppm to 58.3/68.4 and 58.3/66.5 ppm, where the two sets of signals indicate either two sequentially different Thr or the same Thr in two different conformations. Similar division of peaks into multiple signals occurred for a number of other resonances as well. For instance, in the HF sample we observe two Leu and two Val environments, one of each set of signals can be associated with random-coil conformation, and the other ones with extended structures. Histidine signals, including sidechain and backbone carbons, were almost completely absent in the spectra of the AF and HF samples. Histidine residues most likely undergo a *μ*s-ms time-scale conformational exchange which broadens their signals beyond detection, and apparently this motion is independent of the hydration level. Intriguingly, the lack of His signals in any solid-state spectra is consistent with the observed elevated *R*_2_ rate in the solution-state relaxation measurement of the H32 backbone ^15^N, suggesting their role as conformational junctions at the interface between the crystalline and amorphous domains in all experimental conditions. The hydration-induced rearrangement from a predominantly *α*-helical to a *β*-strand-rich structure is also apparent in the ^1^H–^13^C HCH 2D spectra, where the ^1^H dimension gives a further layer of information about the local chemical environments (Fig. 6a–c). Slices taken at the Ala C*α*, Gly C*α*, and Pro C*β* chemical shifts in the HCH spectra of the three samples reveal how the H*α* and H*β* chemical shifts change, in both position and intensity, for these particular residues (Fig. 6d). These changes corroborate the observed shifts of the ^13^C resonances, in particular, the majority of the Ala H*α* resonances shifts from 4.0 ppm (AF) to 5.3 ppm (HF), and a smaller fraction remains around 4.0 ppm, indicating the restructuring of *α*-helix to extended conformations in the ordered segment. The ^1^H dimension also uncovers that two Gly H*α* environments are present in the AP and AF forms with ^1^H shifts at ~4.2 ppm and ~3.1 ppm, while the Gly H*α* shifts merge into one peak at ~3.7 ppm in the HF sample. This latter ^1^H shift is the same that we observed for the Gly H*α* in DMSO solution. From the ^1^H slices it becomes apparent that the Pro side-chains gained much mobility upon hydration which is apparent from the substantial narrowing of the H*β* resonances from 760 Hz in AP, to 260 Hz in AF, all the way to 110 Hz in the HF sample.

**Figure 3.**
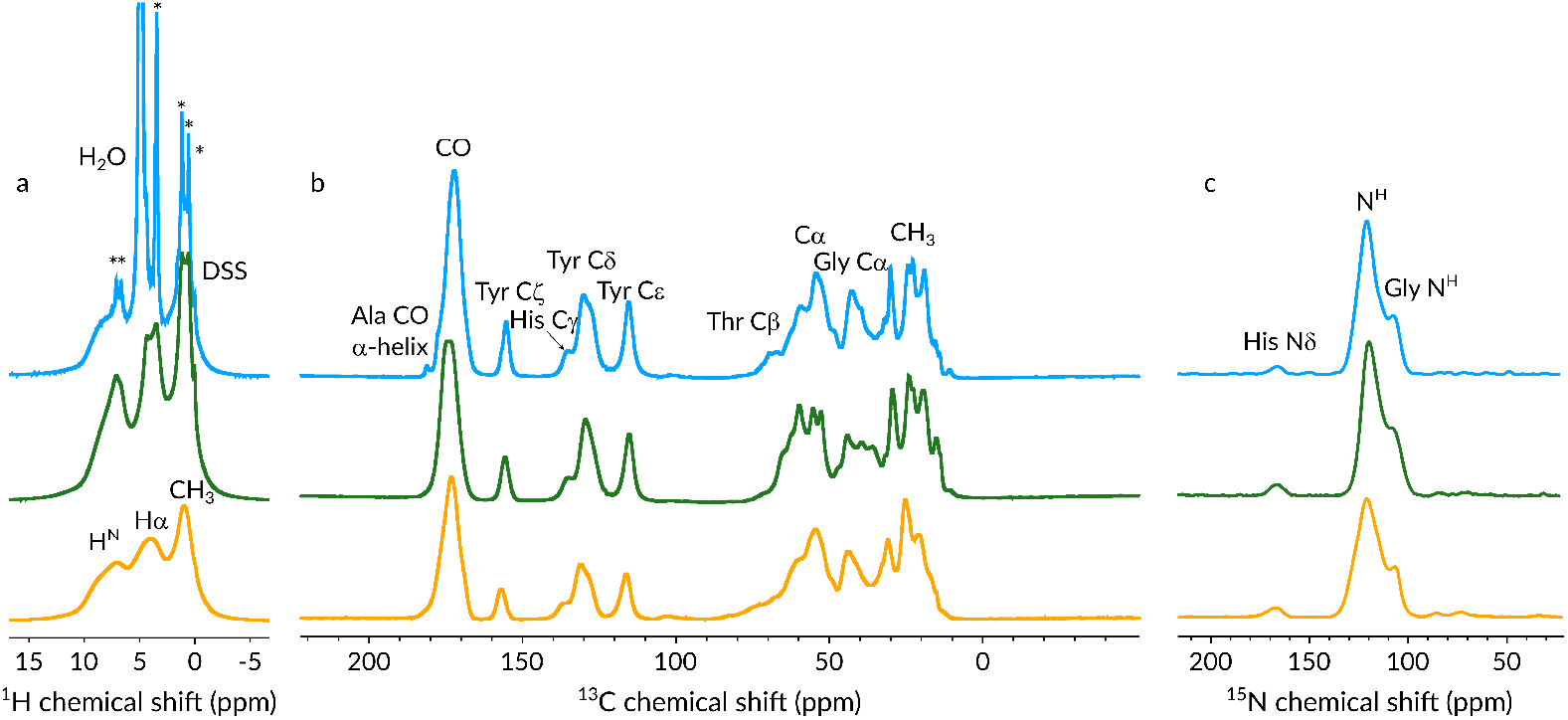
1D ^1^H (a), ^13^C (b), and ^15^N (c) spectra of the AP (orange), AF (green), and HF (blue) TR(1,11) samples measured at 700 MHz ^1^H Larmor frequency at 55.55 kHz MAS. The ^1^H and ^13^C spectra were recorded with direct excitation and detection, the recycle delay was 1.5 and 25 s, respectively. The ^15^N spectra are the ^15^N projections of the ^1^H-detected ^1^H-^15^N 2D CP spectra recorded wit 160 *μ*s CP contact time. Asterisks label the ^1^H resonances of the detergent Triton-X. All spectra are referenced to DSS.

**Figure 4.**
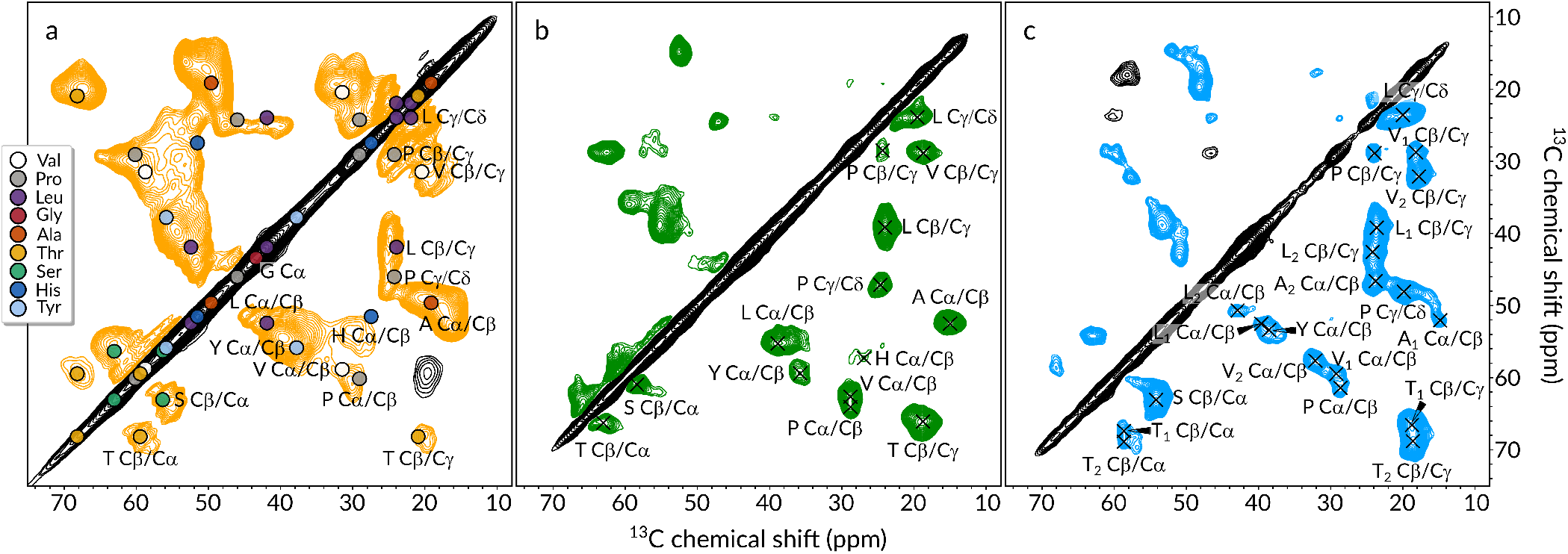
Excerpts from the ^13^C-^13^C DREAM spectra of sample AP (a), AF (b) and HF (c) of TR(1,11) recorded at 700 MHz ^1^H Larmor frequency at 55.55 kHz MAS. In (a) the solution-state ^13^C shifts are labeled on the spectrum color-coded by their amino acid types according to the legend.

**Figure 5.**
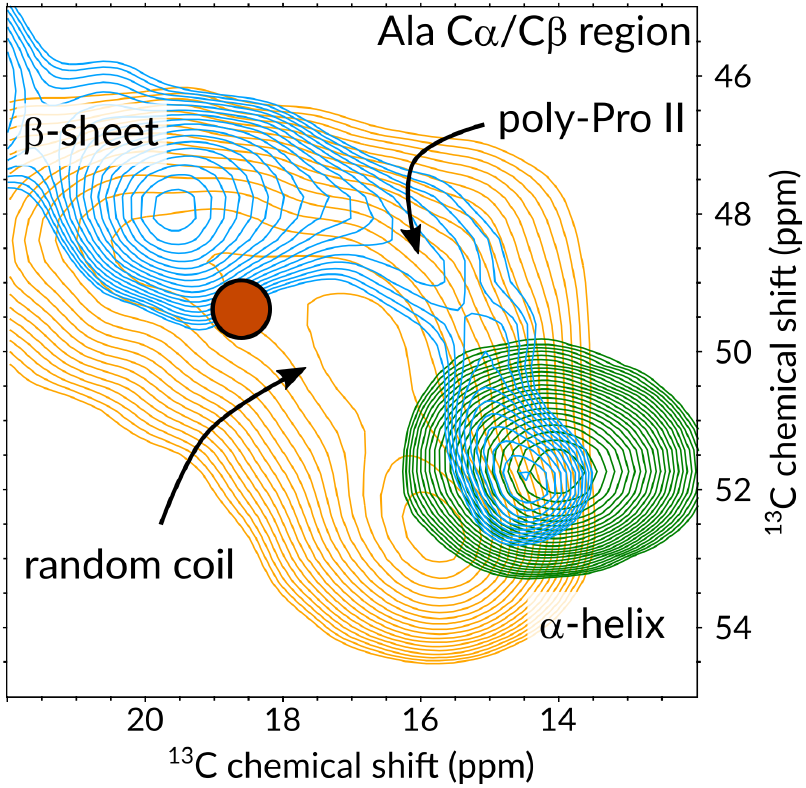
Expanded view of the Ala C*α*/C*β* region showing the typical chemical shift regions for *α*-helical, *β*-sheet, random coil, and poly-proline II helix conformations. Orange, green, and blue spectra are from the AP, AF, and HF samples, respectively. The red dot indicates the average Ala C*α*/C*β* shifts observed in the solution state.

**Figure 6.**
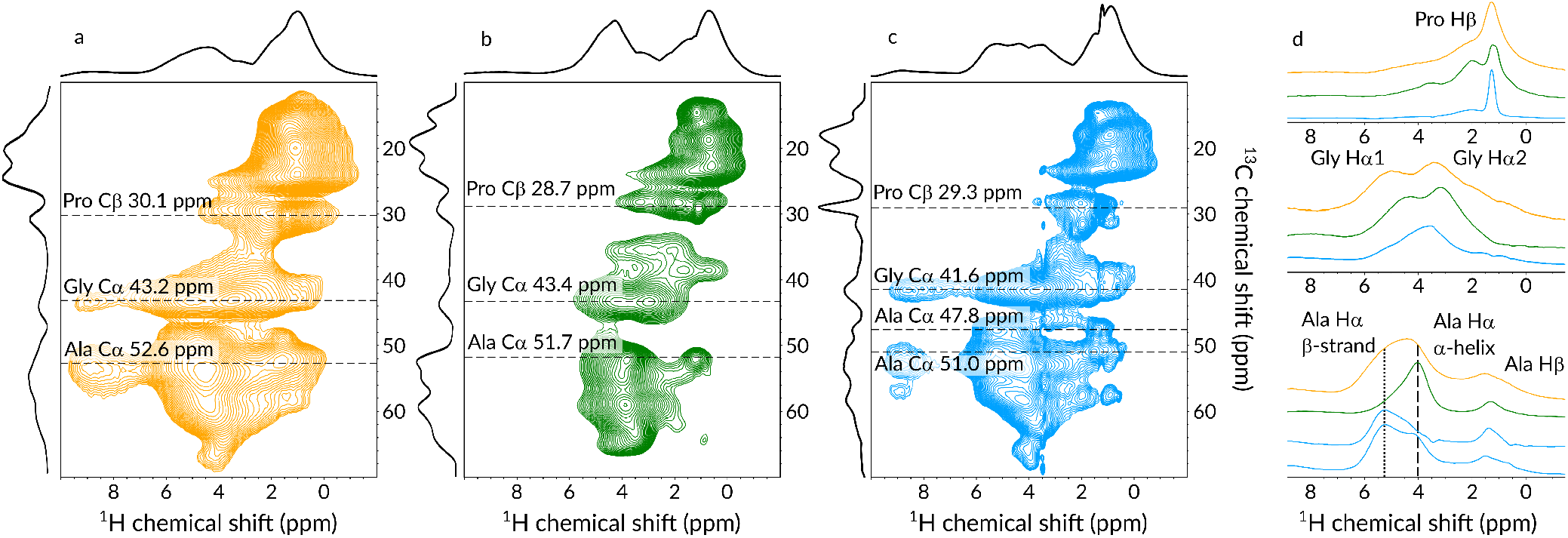
The aliphatic region of the 2D ^1^H-^13^C correlation spectra of the powder (a), ambient film (b), and hydrated film (c) of TR(1,11) measured at 700 MHz ^1^H Larmor frequency with 55.55 kHz MAS frequency. (d) Slices taken at the chemical shift values for Ala C*α*, Gly C*α*, and Pro C*β* are color coded according to the type of the material. Vertical lines indicate the average chemical shift values of Ala H*α* in *β*-strand or *α*-helical conformations.

**Table 1.** ^13^C chemical shift of TR(1,11) compared to literature values. The random coil, *β*-strand and *α*-helix chemical shifts (labeled as “PACSY database”) are reported as the statistical average for 3000+ folded proteins. Reference values are also given for spider (*N. clavipes*) and silkworm (*B. mori*) silks (“Natural silks”). For all Pro entries, values for the *trans* isomer are reported. All shifts are directly referenced to DSS unless labeled with the ^⋆^symbol, where the TMS → DSS conversion was achieved via addition of 1.7 ppm. ^31^ Symbols “-” and “/” indicate a chemical shift range or multiple distinct environments, respectively. An extended version of the table (Table S2) is available in the Supporting Information.

| Residue | PACSY database<sup>29</sup> |  |  | Natural silks |  | Experimental TR(1,11) values |  |  |
| --- | --- | --- | --- | --- | --- | --- | --- | --- |
| | Coil | β-strand | α-helix | Dragline*<sup>31,32</sup> | *B. mori* * <sup>33</sup> | DMSO-*d*<sub>6</sub> | AF | HF |
| Ala CO | 176.9 | 175.6 | 179.1 | 174.3 | 173.9 | 173.5 – 173.8 | 176.8 | 172.0/175.5 |
| Ala Cα | 52.4 | 50.9 | 54.7 | 49.9 | 50.6 | 49.5 – 49.6 | 51.7 | 47.8/51.8 |
| Ala Cβ | 19.4 | 21.7 | 18.5 | 22.6 | 18.3/21.3 | 19.1 – 19.3 | 14.0 | 14.4/19.5 |
| Gly CO | 173.9 | 172.1 | 175.4 | 172.7 | 171.0 | 170.1 – 170.6 | 170.0 – 172.3 | 168.2/170.3 |
| Gly Cα | 45.4 | 45.1 | 46.9 | 45.0 | 44.2 | 43.1 – 43.4 | 42.2 – 45.0 | 41.6 |
| Pro CO | 176.5 | 176.0 | 178.0 | 176.5 | — | 172.8/173.7 | 173.1 | 173.9 |
| Pro Cα | 63.0 | 62.7 | 65.3 | 62.0 | — | 60.5 | 63.4 | 61.3 |
| Pro Cβ | 32.0 | 32.1 | 31.6 | 32.2 | — | 30.1 | 28.7 | 29.3 |
| Pro Cγ | 27.3 | — | — | 27.1 | — | 25.1 | 23.8 | 23.7 |
| Pro Cδ | 50.3 | — | — | 49.2 | — | 47.0 | 46.1 | 46.3 |
| Ser CO | 174.4 | 173.5 | 176.0 | — | 173.9 | 171.4 – 171.9 | 173.1 | 173.9 |
| Ser Cα | 58.2 | 57.1 | 61.0 | 57.2 | 56.3 | 56.3 – 56.8 | 59.0 | 53.7 |
| Ser Cβ | 63.5 | 64.8 | 62.5 | 63.3 | 65.6 | 62.7 – 63.0 | 59.3 | 62.8 |

### Water-induced mobility

After gaining insights into the structural features of the rigid segment of the films, we assessed the highly mobile regions using INEPT-based, solution-state-like 2D experiments. Fig. 7 and S8, S9 compare the ^1^H-detected CP-based and INEPT-based ^1^H–^13^C correlation spectra of the three sample preparations. In the HSQC spectra of both AP and AF, we observed only a handful of peaks in the methyl/aliphatic region (Fig. S9a, b). Upon hydration, multiple new peaks emerged signaling the motional freedom that the water-exposed parts experience (Fig. S8, S9c). The HSQC signals sharpened to an even higher extent, and several additional peaks appeared when the temperature was raised from ~20 to ~40 °C. Sharp signals in the HSQC spectrum of HF with ^1^H line widths of ~50 Hz indicate free, unrestricted motion, comparable to the solubilized form of TR(1,11), although the line widths in DMSO were still an order of magnitude narrower (~5 Hz). The dissolved, highly mobile fragments seem to belong to a separate phase and not to the mobile groups of the rigid core; the Ala, Val, and Leu methyl resonances of the rigid phase appeared as weak and broad signals at the same chemical shift as in the CP-based spectra, while new sets of intense peaks emerged for Ala, Val/Leu, and multiple versions of Pro. We assigned these ^13^C resonances based on their homonuclear ^1^*J*_CC_ coupling patterns, where doublets, triplets, and quartets indicate one, two, or three direct ^13^C covalent bonds. Based on their unique C*β*, C*γ* chemical shifts, we could differentiate between Pro with both cis and trans XaaPro peptide-bond conformations indicating a random-coil-like behavior. The reason behind such intense Pro peaks in the HSQC spectrum is the fast ring puckering which has a timescale of > 10^8^s^-1^in solution state and have been observed to occur on a similar time-scale for prolines in hydrated collagen,^28,34^ elastin,^35^ or MaSp2 silk of A. aurantia.^36^ Using the ^13^C multiplicity we could assign a few other peaks as belonging to CH_3_of Ala (or Val/Leu), or to CH groups of Val or Leu, and we also observed a weak Gly H*α*/C*α* correlation at 3.04/40.2 ppm (Fig. 7 right). The assignment was further facilitated by INEPT-transfer-based ^13^C–^13^C correlation spectrum using through-space RFDR mixing (Fig. 7 left). Strong exchange cross-peaks between the P_trans_ and P_cis_ C*β* resonances reveal a slow conformational exchange process between the two isomers. Besides, we see strong cross-peaks between P_cis_ C*γ* (21.8 ppm) and a methyl group (12.8 ppm), and another one between Ptrans C*β* (27.9 ppm) and another methyl group (14.7 ppm). Based on the sequential proximity of P22 and A23, we assigned both of these methyl resonances to A23C*β* carbons and thus we concluded that the GPA segment is the one that is most affected by hydration. The upfield shifted Ala C*β* chemical shifts are indicative of a helical backbone conformation. In this aqueous phase, we do not see any evidence for alanines having extended, *β*-stranded structures which is contrary to what we have observed in DMSO-*d*_6_ solution, implying that water genuinely interacts with the peptide stretch.

**Figure 7.**
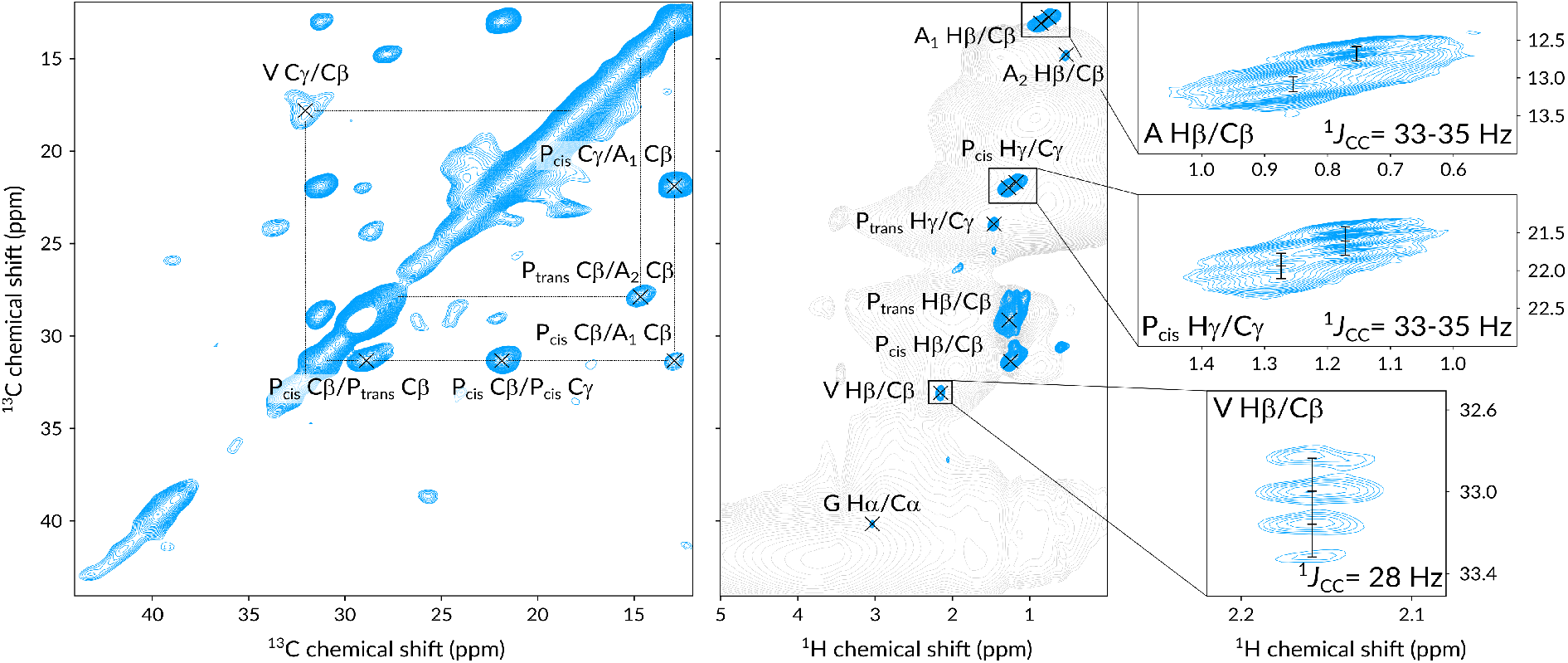
Refocused-INEPT-based ^13^C–^13^C RFDR (left) and ^1^H–^13^C HSQC (middle) spectra of HF, measured at ~20 °C, and ~40 °C, respectively. Gray spectrum marks the silhouette of the CP-based ^1^H–^13^C correlation spectrum. Insets show the same HSQC spectrum as in the middle but recorded with 43 ms indirect acquisition time with examples for doublet (top), triplet (center), and quartet (bottom) fine structures assigned to Ala C*β*, Procis C*γ*, and Val C*β*.

## Discussion

The atomistic insight provided by solution and solid-state NMR is key to understand the thermo-mechanical and physico-chemical properties of tandem repeat biomaterials. Previous studies on TR proteins have established that films made from TR(1, *n*) proteins with increasing repetition number rapidly and reversibly switch between states with remarkably different thermal conductivity (*κ*) when the film is hydrated or the water is removed.^10^ Such exceptional thermal transport was attributed to relative changes in specific heat, phonon velocity as well as phonon scattering distance which is correlated to mean square deviations of hydrogens. Our study elaborates on the structural basis of this hydration-dependent property. Based on the solidstate NMR data we are confident that the two factors that contribute to the increased κ are the restructuring of the protein matrix into a conformation that is rich in *β*-strands and the increased overall nanosecond time-scale mobility thanks to the plasticizing effect of water. In this respect, higher structural order, in particular high degree of ordered *β*-pleated sheets, favors phonon energy transport, while the increased overall dynamics lead to higher molar diffusivity. Our data suggests that it is not the amorphous region which gains the highest mobility upon hydration, but rather the GPA triplet, where this interaction results in *cis-trans* isomerization of the GP peptide bond and in fast proline ring pucker fluctuations. In collagen, similar fast ring puckering in spatially clustered glycine-proline-hydroxyproline triplets contribute to low energy compression-extension and bending of the triple helix.^37^ We propose that the GPA sequence in the hydrated TR(1,11) film has a related role as it provides high conformational flexibility at the junction between the Gly-rich and Ala-rich regions, which in turn leads to a protein network with increased elasticity and toughness.^9^ We also observed that the Gly-rich segments are partially converted into rigid *β*-strands which are likely to enhance the overall crystallinity and hence contribute to the thermal conductivity of the material. Similar random-coil-to-*β*-sheet transformation in the Gly-rich regions was reported for silk fibroin fibers of *Samia cynthia ricini* and *Bombyx mori* when the fibers where stretched.^38^ Likewise, nearly 30% fraction of the glycines in the water plasticized state of the dragline silk of *Nephila clavipes* were found in extended conformation highlighting the multifaceted role of Gly in the amorphous segments of tandem repeat proteins.^39^ Hydration initiates large-scale structural rearrangements in the ordered, rigid domain as well. Since the ambient film was cast from an HFIP solution, therefore the strong helix-inducing property of the fluorinated solvent resulted in a film that was also rich in *α*-helices; a trait found in other protein films cast from HFIP.^40,41^ Upon hydration, most of the helical parts were transformed into extended *β*-strands, and only ~10% of the alanines remained in helical conformation. Hydration-induced conversion between *α*-helical and *β*-strand-rich conformations have been observed in the ordered domain of both natural and artificial silks and amyloidogenic peptides where helical conformation is preferred in hydrophobic while *β*-strands in hydrophilic environments.^42–44^ Among all the amino acids of TR(1,11), the two prolines at the junction between the Ala-rich and Gly-rich segments emerged as the main drivers in the formation of secondary structures and in defining the molecule’s dynamic properties. In the solution state, we observed a clear *β*-strand propensity initiated by the Pro preceding the Ala-rich segment. The Pro at the end of the crystalline unit, as part of the VHHP segment, was found to be involved in slow time-scale conformational exchange facilitated by the motion of the preceding His. These structural and motional features are mirrored in the hydrated solid sample, where the lack of His signals suggest a large amplitude *μ*s-ms time-scale fraying motion at the end of the *β*-strands. The motional freedom of His side-chains likely contributes to the remarkable proton conductivity of the hydrated TR films which is associated with fluctuations happening on the very same time scale.^11^ Lastly, we propose that the dramatic increase in *β*-strand content of the hydrated film plays a major role in determining the outstanding self-healing capabilities of TR(1,11). Molecular dynamics simulations have indicated that the YGYGGLYGGLYGGLGY peptide sequence is responsible for the autohesion phenomenon occurring in the rubbery state.^12^ It has also been shown that hydration lowers the glass transition temperature, enabling a stable rubbery-state over a wide range of temperatures.^11^ It was proposed that—in the water-plasticized rubbery state—the Gly-rich amorphous segments of SRT proteins align along the strain direction at low stress.^16^ Our data suggest that such alignment is facilitated by the high *β*-strand content gained upon hydration which drives the self-healing mechanism on the microscopic scale and permits the self-adhesion of SRT-based thermoplastic polymers.

## Conclusions

In this study, we investigated the conformation and dynamics of a recombinant tandem repeat protein TR(1,11) inspired by the amino acid sequence of the squid ring teeth protein of Loligo vulgaris. Using advanced NMR spectroscopy methods, we observed that the mainly disordered TR(1,11) in DMSO solution shows a clear tendency towards extended structures. Film casting from HFIP solution induces extensive *α*-helix formation that, upon hydration, transforms into a *β*-strand-rich protein matrix where strong and cooperative H-bonds stabilizes the nanoconfined *β*-sheets, in the formation of which both the Ala-rich and the Gly-rich segments contribute. Water interacts directly and indirectly with the protein film; it directly solubilizes the junction between the amorphous and ordered domains by elevating the mobility of the proline and alanine side-chains at the water exposed GPA sequence; while indirectly affects the overall structure by inducing a large-scale structural rearrangement in the ordered rigid domain. In summary, we demonstrate how the aforementioned structural features define the origin of the striking macroscopic characteristics of TR(1,11), namely elasticity, thermal conductivity, and self-healing capability. Our study provides insights into the mechanistic and residue-specific behavior of the protein and assists in the rational design of *de novo* proteinaceous materials.

## Acknowledgments

P.R. acknowledges support from the Deutsche Forschungsgemeinschaft (DFG, German Research Foundation) – SFB 1309 – 325871075, the Center for NanoScience (CeNS), the Fonds der Chemischen Industrie, and Universitätsgesellschaft München. M.C.D. and H.J. were supported by the United States Army Research Office (grant no. W911NF-16-1-0019 and W911NF-18-1-026) and the Huck Endowment of The Pennsylvania State University. We thank Thea Schinkel and Pia Heinrichs for critical reading of the manuscript.

## Materials and methods

### Gene editing

Repetitive DNA constructs was described in our previous work.^8^ Briefly, unit repeating gene fragments (*n* = 1) originated from European squid tentacles (*Loligo vulgaris*) including cloning sequences were purchased from Genewiz (South Plainfield, NJ, USA). Rolling circle amplification was performed by BbvCI and phi29-polymerase, and protected digestion was performed by Acc65I and ScaI. Near 1250 bp regions of double-stranded products of protected digestion of rolling circle amplification were inserted into modified pET14b backbones were selected by agarose-gel-cutting and purified by Omega Bio-Tek E.Z.N.A. gel extraction kit. Purified templates were cloned into the open-reading frame of pET14b vector. Sequence verified plasmids (by Sanger-sequencing, Genomic core facility at Pennsylvania State University) were transformed into the BL21 (DE3) expression strain for expression. Subsequently, the recombinant *E. coli* strains were grown on Luria-Bertani (LB) agar plates supplemented with carbenicillin (100 g/L) and incubated at 37 °C for 13 - 15 hours.

### Protein expression and extraction

All isotope components were purchased from Cambridge Isotope Laboratories, Inc. (Tewksbury, MA, USA) and LB broth (Miller) was purchased from Fisher Scientific (Waltham, MA, USA). 50 mL of LB broth (1.25 g of LB broth powder per 50 mL of double distilled water) was used as a starter culture and 500 ml of M9 minimal media with ^15^NH_4_Cl and ^13^C– glucose for each protein expression. A single colony from the plate of expression strains was used to inoculate 50 ml of LB-Miller incubating at 29 °C and 210 rpm for about 10 hours as a starter culture (OD_600_ =~1, pH = ~7). 10 mL of fully-grown starter culture was added to 0.5 L of M9 media and incubated at 37 °C and 300 rpm for 5 - 7 hours. 0.5 L of M9 media with filter sterilized isopropyl *β*-D-1-thiogalactopyranoside (IPTG) of 1 mM as a final concentration for 8 hours, and then harvested by centrifugation (10000 rpm for 15 minutes). Cell lysis was performed by mechanical method by a high-pressure homogenizer (Microfluidizer, Microfluidics-M110EH-30) at 1034 bar, and pellets were collected by centrifugation with 10000 rpm for 30 minutes. Protein isolation was performed by low-spin centrifugation. Lysed pellets were washed with 50 mL of urea-extraction buffer (100 mM Tris pH 7.4, 5 mM EDTA, 2 M Urea, 2% (v/v) Triton X-100, one time) and 50 mL last-extraction buffer (100 mM Tris, pH 7.4, 5 mM EDTA, two times) to remove left-over cell debris generated during physical extraction process by centrifugation (at 5000 rpm). Washed pellets were lyophilized with FreeZone 6 plus (Lab-conco) for 12 hours at CSL Behring fermentation facility at Pennsylvania State University. The yield of protein expression was approximately 10-20 mg per 500 mL of M9 minimal medium.

### Protein film casting

The lyophilized purified protein was completely dissolved in hexafluoroisopropanol (HFIP) to a final concentration of 20 mg/mL. Each film was cast by pouring HFIP-protein solutions to PDMS molds. Average mass of cast films was ~5 mg.

### ATR-FTIR

Attenuated total reflectance Fourier transform infrared spectral data were collected (Thermo Scientific Nicolet 6700 FT-IR) under attenuated total reflection (diamond crystal) mode using Happ-Genzel apodization with 4 cm^-1^resolution from 400 to 4000 cm^-1^. For each spectrum 256 scans were co-added. Gaussian curve fitting was performed in OriginPro 8.5 software by using a nonlinear least-squares method.

### Solution-state NMR

All solution-state NMR experiments were performed at 37 °C on a Bruker Avance III spectrometer operating at a ^1^H Larmor frequency of 800 MHz (corresponding to a magnetic field of 18.79 T) equipped with a triple channel, cryogenically cooled HCN TCI probe. ^1^H, ^13^C, and ^15^N shifts were calibrated to DSS based on IUPAC recommendations using an internal standard.^45^ Triple resonance assignment (HNCA, HNCO, HNCACB and hCC-coNH) experiments were recorded employing Band-selective Excitation Short-Transient (BEST) techniques and analyzed using Cara.^46,47^ ^15^N *R*_1_rates were obtained with 8 relaxation delays of 20, 50, 100, 150, 200, 300, 400, and 500 ms. ^15^N *R*_2_ rates were measured via a standard Carr-Purcell-Meiboom-Gill (CPMG) pulse sequence. The measurement was set up with 12 linearly incremented CPMG pulse trains of the duration of 16.96 ms each. Heteronuclear nOe data was obtained via direct comparison of a spectrum recorded with a 3 s presaturation pulse on the proton channel, and a reference spectrum without presaturation. In both cases, relaxation delay was set to 5 s. All spin relaxation data sets were collected via ^1^H–^15^N correlated HSQC spectra recorded in an interleaved fashion. The extraction of peak intensities and signal-to-noise ratios was done with NMRFAM-Sparky.^48^ Relaxation rates were fitted with a two-parameter monoexponentially decaying function. The uncertainties of the relaxation rates were propagated to the errors of the spectral densities using standard error propagation rules. The relaxation rates were analyzed with the reduced spectral density mapping (RSDM) formalism.^25^ ^1^H DOSY experiments were measured in a pseudo-2D fashion with Δ = 150 ms and *δ* = 2 ms and 64 increments using a standard pulse sequence.

### Solid-state NMR

All solid-state NMR experiments were recorded using a Bruker Neo NMR spectrometer operating at a ^1^H Larmor frequency of 700.4 MHz equipped with a triple-resonance HCN 1.3 mm MAS probe. The MAS frequency was set to 55.55 kHz and it was stable within 10 Hz. Unless otherwise stated, the temperature was regulated using a VT gas flow to 0 °C which corresponds to a ~20 °C nominal temperature within the rotor assessed by the water resonance frequency with respect to internal DSS signal. Typical experimental parameters were 1.5 *μ*s, 2.7 *μ*s, and 3.5 *μ*s 90° pulse length for ^1^H, ^13^C, and ^15^N, respectively. In the ^1^H–^13^C CP-based experiments, the ^1^H field strength was ramped with a tangential shape with an effective strength of 68 kHz and a 10 kHz rectangular shape was applied on the ^13^C channel with a contact time of 1 – 2 ms. The ^1^H–^15^N CP step involved a 12 kHz tangential ^1^H shape and a 42 kHz rf field on ^15^N, the contact time was 160 *μ*s. A 12.5 kHz XiX ^1^H-decoupling was applied during the ^13^C or ^15^N acquisition, and a 10 kHz WALTZ-16 decoupling was used during ^1^H detection. Water signal in the hydrated sample was suppressed by the MISSISSIPPI pulse sequence with 15 kHz irradiation for 80 ms. The INEPT transfer delay (1/4J) was 2 ms (^1^H–^13^C). All ^1^H–^13^C or ^1^H–^15^N 2D experiments were recorded in a ^1^H-detected fashion with a recycle delay of 1 – 2 s. Homonuclear ^13^C–^13^C mixing was achieved with either dipolar recoupling enhanced by amplitude modulation (DREAM) or with radio-frequency-driven recoupling (RFDR) approaches. The ^13^C–^13^C DREAM mixing time was 4 ms, and a tangential shaped pulse was centered at 56 ppm, the ^13^C–^13^C RFDR mixing time was 4.6 ms with the carrier placed at 24 ppm. ^49^In the DREAM experiments the spectral width of the indirect dimension was 55555 Hz, with a maximum *t*_1_ evolution time of 5.4 ms over 600 increments, recorded with 64 scans. In the refocused-INEPT-based ^13^C–^13^C 2D experiment the spectral width of the indirect dimension was 27777 Hz, with a maximum *t*_1_ evolution time of 10.8 ms over 600 increments, recorded with 32 scans. In the ^1^H–^13^C CP-based 2D HCH experiment the spectral width of the indirect dimension was 27777 Hz, with a maximum *t*_1_ evolution time of 3.6 ms over 200 increments, recorded with 128 scans; the ^13^C transmitter frequency was set to either 50 ppm or 173 ppm for measuring the aliphatic or the aromatic and carbonyl region of the HCH spectrum, respectively. In the ^1^H–^13^C INEPT-based HSQC experiment the spectral width of the indirect dimension was 27777 Hz, with a maximum *t*_1_ evolution time of 10.8 ms over 600 increments, recorded with 96 scans, or with a maximum *t*_1_ evolution time of 43.2 ms over 2400 increments, recorded with 16 scans, in both experiments the ^13^C carrier was placed to 50 ppm. In the ^1^H–^15^N 2D HNH experiment the spectral width of the indirect dimension was 13888 Hz, with a maximum *t*_1_ evolution time of 4.3 ms over 120 increments, recorded with 256 scans with ^15^N transmitter frequency set to 120 ppm. ^1^H, ^13^C, and ^15^N chemical shifts in all solid state spectra were referenced to internal DSS signal. The protein film was hydrated by soaking the opened rotor in H_2_O overnight at 4 °C.

## References

(1) Altman, G. H.; Diaz, F.; Jakuba, C.; Calabro, T.; Horan, R. L.; Chen, J.; Lu, H.; Richmond, J.; Kaplan, D. L. Silk based biomaterials. Biomaterials 2003, 39, 922–931.

(2) Almine, J. F.; Bax, D. V.; Mithieux, S. M.; Smith, L. N.; Rnjak, J.; Waterhouse, A.; Wise, S. G.; Weiss, A. S. Elastin-based materials. Chem. Soc. Rev. 2010, 39, 3371–3379.

(3) Sun, J.; Su, J.; Ma, C.; Göstl, R.; Herrmann, A.; Liu, K.; Zhang, H. Fabrication and Mechanical Properties of Engineered Protein-Based Adhesives and Fibers. Adv. Mater. 2020, 32, 1–16.

(4) Sutherland, T. D.; Huson, M. G.; Rapson, T. D. Rational design of new materials using recombinant structural proteins: Current state and future challenges. J. Struct. Biol. 2018, 201, 76–83.

(5) Pena-Francesch, A.; Demirel, M. C. Squid-inspired tandem repeat proteins: Functional fibers and films. Front. Chem. 2019, 7, 1–16.

(6) Guerette, P. A.; Hoon, S.; Ding, D.; Amini, S.; Masic, A.; Ravi, V.; Venkatesh, B.; Weaver, J. C.; Miserez, A. Nanoconfined *β*-sheets mechanically reinforce the supra-biomolecular network of robust squid Sucker Ring Teeth. ACS Nano 2014, 8, 7170–7179.

(7) Pena-Francesch, A.; Florez, S.; Jung, H.; Sebastian, A.; Albert, I.; Curtis, W.; Demirel, M. C. Materials fabrication from native and recombinant thermoplastic squid proteins. Adv. Funct. Mater. 2014, 24, 7401–7409.

(8) Jung, H.; Pena-Francesch, A.; Saadat, A.; Sebastian, A.; Kim, D. H.; Hamilton, R. F.; Albert, I.; Allen, B. D.; Demirel, M. C. Molecular tandem repeat strategy for elucidating mechanical properties of high-strength proteins. Proc. Natl. Acad. Sci. U. S. A. 2016, 113, 6478–6483.

(9) Pena-Francesch, A.; Jung, H.; Segad, M.; Colby, R. H.; Allen, B. D.; Demirel, M. C. Mechanical Properties of Tandem-Repeat Proteins Are Governed by Network Defects. ACS Biomater. Sci. Eng. 2018, 4, 884–891.

(10) Tomko, J. A.; Pena-Francesch, A.; Jung, H.; Tyagi, M.; Allen, B. D.; Demirel, M. C.; Hopkins, P. E. Tunable thermal transport and reversible thermal conductivity switching in topologically networked bio-inspired materials. Nat. Nanotechnol. 2018, 13, 959–964.

(11) Pena-Francesch, A.; Jung, H.; Hickner, M. A.; Tyagi, M.; Allen, B. D.; Demirel, M. C. Programmable Proton Conduction in Stretchable and Self-Healing Proteins. Chem. Mater. 2018, 30, 898–905.

(12) Sariola, V.; Pena-Francesch, A.; Jung, H.; Cetinkaya, M.; Pacheco, C.; Sitti, M.; Demirel, M. C. Segmented molecular design of self-healing proteinaceous materials. Sci. Rep. 2015, 5, 1–9.

(13) Pena-Francesch, A.; Jung, H.; Demirel, M. C.; Sitti, M. Biosynthetic self-healing materials for soft machines. Nat. Mater. 2020, 19.

(14) Yllmaz, H.; Pena-Francesch, A.; Shreiner, R.; Jung, H.; Belay, Z.; Demirel, M. C.; Özdemir, K.; Yang, L. Structural protein-based whispering gallery mode resonators. ACS Photonics 2017, 4, 2179–2186.

(15) Sang, Y.; Li, M.; Liu, J.; Yao, Y.; Ding, Z.; Wang, L.; Xiao, L.; Lu, Q.; Fu, X.; Kaplan, D. L. Biomimetic Silk Scaffolds with an Amorphous Structure for Soft Tissue Engineering. ACS Appl. Mater. Interfaces 2018, 10, 9290–9300.

(16) Rieu, C.; Bertinetti, L.; Schuetz, R.; Salinas-Zavala, C. C.; Weaver, J. C.; Fratzl, P.; Miserez, A.; Masic, A. The role of water on the structure and mechanical properties of a thermoplastic natural block co-polymer from squid sucker ring teeth. Bioinspiration and Biomimetics 2016, 11.

(17) Goldberga, I.; Li, R.; Duer, M. J. Collagen Structure-Function Relationships from Solid-State NMR Spectroscopy. Acc. Chem. Res. 2018, 51, 1621–1629.

(18) Ghosh, M.; Prajapati, B. P.; Kango, N.; Dey, K. K. A comprehensive and comparative study of the internal structure and dynamics of natural *β*-keratin and regenerated*β*-keratin by solid state NMR spectroscopy. Solid State Nucl. Magn. Reson. 2019, 101, 1–11.

(19) Yao, X. L.; Hong, M. Structure Distribution in an Elastin-Mimetic Peptide (VPGVG)3 Investigated by Solid-State NMR. J. Am. Chem. Soc. 2004, 126, 4199–4210.

(20) Yu, P. Synchrotron IR microspectroscopy for protein structure analysis: Potential and questions. Spectroscopy 2006, 20, 229–251.

(21) Alderson, T. R.; Benesch, J. L.; Baldwin, A. J. Proline isomerization in the C-terminal region of HSP27. Cell Stress Chaperones 2017, 22, 639–651.

(22) Kawagoe, S.; Nakagawa, H.; Kumeta, H.; Ishimori, K.; Saio, T. Structural insight into proline cis/trans isomerization of unfolded proteins catalyzed by the trigger factor chaperone. J. Biol. Chem. 2018, 293, 15095–15106.

(23) Aulikki, M.; Ravotti, F.; Arai, H.; Glabe, C. G.; Wall, J. S.; Böckmann, A. Atomic-resolution structure of a disease-relevant. Proc. Natl. Acad. Sci. U. S. A. 2016, 113, 4976–4984.

(24) Wilkins, D. K.; Grimshaw, S. B.; Receveur, V.; Dobson, C. M.; Jones, J. A.; Smith, L. J. Hydrodynamic radii of native and denatured proteins measured by pulse field gradient NMR techniques. Biochemistry 1999, 38, 16424–16431.

(25) Peng, J. W.; Wagner, G. Mapping of the Spectral Densities of N-H Bond Motions in Eglin C Using Heteronuclear Relaxation Experiments. Biochemistry 1992, 31, 8571–8586.

(26) Daragan, V. A.; Mayo, K. H. Motional model analyses of protein and peptide dynamics using 13C and 15N NMR relaxation. Prog. Nucl. Magn. Reson. Spectrosc. 1997, 31, 63–105.

(27) Yao, X. L.; Conticello, V. P.; Hong, M. Investigation of the dynamics of an elastin-mimetic polypeptide using solid-state NMR. Magn. Reson. Chem. 2004, 42, 267–275.

(28) Saitô, H.; Yokoi, M. A 13C NMR study on collagens in the solid state: Hydration/ dehydration-induced conformational change of collagen and detection of internal motions. J. Biochem. 1992, 111, 376–382.

(29) Fritzsching, K. J.; Hong, M.; Schmidt-Rohr, K. Conformationally selective multidimensional chemical shift ranges in proteins from a PACSY database purged using intrinsic quality criteria. J. Biomol. NMR 2016, 64, 115–130.

(30) Holland, G. P.; Creager, M. S.; Jenkins, J. E.; Lewis, R. V.; Yarger, J. L. Determining secondary structure in spider dragline silk by carbon-carbon correlation solid-state NMR spectroscopy. J. Am. Chem. Soc. 2008, 130, 9871–9877.

(31) Jenkins, J. E.; Holland, G. P.; Yarger, J. L. High resolution magic angle spinning NMR investigation of silk protein structure within major ampullate glands of orb weaving spiders. Soft Matter 2012, 8, 1947–1954.

(32) Jenkins, J. E.; Creager, M. S.; Butler, E. B.; Lewis, R. V.; Yarger, J. L.; Holland, G. P. Solid-state NMR evidence for elastin-like *β*-turn structure in spider dragline silk. Chem. Commun. 2010, 46, 6714–6716.

(33) Asakura, T.; Sugino, R.; Yao, J.; Takashima, H.; Kishore, R. Comparative structure analysis of tyrosine and valine residues in unprocessed silk fibroin (silk I) and in the processed silk fiber (silk II) from Bombyx mori using solid-state 13C,15N, and 2H NMR. Biochemistry 2002, 41, 4415–4424.

(34) London, R. E. On the Interpretation of 13C Spin-Lattice Relaxation Resulting from Ring Puckering in Proline. J. Am. Chem. Soc. 1978, 100, 2678–2685.

(35) Dabalos, C. L.; Ohgo, K.; Kumashiro, K. K. Detection of Labile Conformations of Elastin’s Prolines by Solid-State Nuclear Magnetic Resonance and Fourier Transform Infrared Techniques. Biochemistry 2019, 58, 3848–3860.

(36) Shi, X.; Yarger, J. L.; Holland, G. P. Elucidating proline dynamics in spider dragline silk fibre using 2H-13C HETCOR MAS NMR. Chem. Commun. 2014, 50, 4856–4859.

(37) Chow, W. Y.; Forman, C. J.; Bihan, D.; Puszkarska, A. M.; Rajan, R.; Reid, D. G.; Slatter, D. A.; Colwell, L. J.; Wales, D. J.; Farndale, R. W.; Duer, M. J. Proline provides site-specific flexibility for in vivo collagen. Sci. Rep. 2018, 8, 1–13.

(38) Asakura, T.; Ito, T. Structure of alanine and glycine residues of samia cynthia ricini silk fibers studied with solid-state 15N and13C NMR. Macromolecules 1999, 32, 4940–4946.

(39) Holland, G. P.; Jenkins, J. E.; Creager, M. S.; Lewis, R. V.; Yarger, J. L. Quantifying the fraction of glycine and alanine in *β*-sheet and helical conformations in spider dragline silk using solid-state NMR. Chem. Commun. 2008, 5568–5570.

(40) Tasei, Y.; Nishimura, A.; Suzuki, Y.; Sato, T. K.; Sugahara, J.; Asakura, T. NMR Investigation about Heterogeneous Structure and Dynamics of Recombinant Spider Silk in the Dry and Hydrated States. Macromolecules 2017, 50, 8117–8128.

(41) Moseti, K. O.; Yoshioka, T.; Kameda, T.; Nakazawa, Y. Structure water-solubility relationship in *α*-helix-rich films cast from aqueous and 1,1,1,3,3,3-hexafluoro-2-propanol solutions of S. c. ricini silk fibroin. Molecules 2019, 24, 12–13.

(42) Kumar, S. T.; Leppert, J.; Bellstedt, P.; Wiedemann, C.; Fändrich, M.; Görlach, M. Solvent Removal Induces a Reversible *β*-to-*α* Switch in Oligomeric A*β* Peptide. J. Mol. Biol. 2016, 428, 268–273.

(43) Otikovs, M.; Andersson, M.; Jia, Q.; Nordling, K.; Meng, Q.; Andreas, L. B.; Pintacuda, G.; Johansson, J.; Rising, A.; Jaudzems, K. Degree of Biomimicry of Artificial Spider Silk Spinning Assessed by NMR Spectroscopy. Angew. Chemie-Int. Ed. 2017, 56, 12571–12575.

(44) Saltô, H.; Tabeta, R.; Shoji, A.; Ozaki, T.; Ando, I. Conformational Characterization of Polypeptides in the Solid State as Viewed from the Conformation-Dependent 13C Chemical Shifts Determined by the 13C Cross Polarization/Magic Angle Spinning Method: Oligo(L-alanine), Poly(L-alanine), Copolymers of L-and. Macromolecules 1983, 16, 1050–1057.

(45) Markley, J. L.; Bax, A.; Arata, Y.; Hilbers, C. W.; Kaptein, R.; Sykes, B. D.; Wright, P. E.; Wüthrich, K. Recommendations for the presentation of NMR structures of proteins and nucleic acids. IUPAC-IUBMB-IUPAB inter-union task group on the standardization of data bases of protein and nucleic acid structures determined by NMR spectroscopy. Eur. J. Biochem. 1998, 256, 1–15.

(46) Keller, R. The computer aided resonance assignment tutorial. Goldau, Switz. Cantina Verlag 2004, 1–81.

(47) Lescop, E.; Schanda, P.; Brutscher, B. A set of BEST triple-resonance experiments for time-optimized protein resonance assignment. J. Magn. Reson. 2007, 187, 163–169.

(48) Lee, W.; Tonelli, M.; Markley, J. L. NMRFAM-SPARKY: Enhanced software for biomolecular NMR spectroscopy. Bioinformatics 2015, 31, 1325–1327.

(49) Verel, R.; Ernst, M.; Meier, B. H. Adiabatic dipolar recoupling in solid-state NMR: The DREAM scheme. J. Magn. Reson. 2001, 150, 81–99.

