## Supplementary Information file for "Hydration-induced structural transitions in biomimetic tandem repeat proteins"

21/12/2020

#### Table of Contents

**1 Supplementary Methods** **S2**
1.1 Protein sequence . . . . . S2
1.2 Hydrodynamic radius calculation . . . . . S2

#### List of Figures

S1 MALDI-TOF spectra of TR(1,11) protein . . . . . S2
S4 $^{1}$H-$^{15}$N CP-based 2D spectra of AP, AF, and HF . . . . . S4
S5 Pulse sequence of the refocused INEPT-based 2D $^{13}$C–$^{13}$C experiment . . . . . S4
S6 $^{1}$H-$^{13}$CO CP-based 2D spectra of sample AP, AF, and HF . . . . . S5
S7 $^{1}$H-$^{13}$CO CP-based 2D spectra of sample AP, AF, and HF . . . . . S5
S8 Excerpt from the aliphatic/methyl region of the $^{1}$H-$^{13}$C HSQC spectrum of HF . . . . . S6
S9 Comparison of the methyl region of the $^{1}$H-$^{13}$C HSQC spectra of AP, AF, and HF . . . . . S6

#### List of Tables

S1 Deconvoluted FTIR spectral data for casted and washed films. . . . . . S7
S2 $^{13}$C chemical shift of TR(1,11) compared to literature values . . . . . S7

### 1 Supplementary Methods

#### 1.1 Protein sequence

MGTLS**YGYGGLYGGLYGGLGYGPAAASVSTVHHP**
STGTLS**YGYGGLYGGLYGGLGYGPAAASVSTVHHP**
STGTLS**YGYGGLYGGLYGGLGYGPAAASVSTVHHP**
STGTLS**YGYGGLYGGLYGGLGYGPAAASVSTVHHP**
STGTLS**YGYGGLYGGLYGGLGYGPAAASVSTVHHP**
STGTLS**YGYGGLYGGLYGGLGYGPAAASVSTVHHP**
STGTLS**YGYGGLYGGLYGGLGYGPAAASVSTVHHP**
STGTLS**YGYGGLYGGLYGGLGYGPAAASVSTVHHP**
STGTLS**YGYGGLYGGLYGGLGYGPAAASVSTVHHP**
STGTLS**YGYGGLYGGLYGGLGYGPAAASVSTVHHP**
STGTLS**YGYGGLYGGLYGGLGYGPAAASVSTVHHP**
STGTLS**YGYGGLYGGLYGGLGYGP**

Black, red and blue segments indicate the cleveage site, the amorphous and the crystalline region, respectively.

#### 1.2 Hydrodynamic radius calculation

For a large solute molecule of radius $R$ in a solvent of much smaller molecules, the diffusion coefficient $D$ of the solute is described via the Stokes-Einstein relationship:

$$R_h = \frac{k_B T}{\pi D \eta \epsilon} \tag{S1}$$

where $R_h$ is the hydrodynamic radius of the molecule, $k_B$ is the Boltzmann constant, $T$ is the temperature of the system, $\eta$ is the solvent viscosity and $\epsilon$ is a constant determined by the choice of stick ($\epsilon = 6$) or slip ($\epsilon = 4$) hydrodynamic boundary conditions at the solute surface. Consistently with previous investigations on polymers of similar molecular mass, we opted for constant $\epsilon = 4$,[1] which resulted in $R_h = 33.0$ Å.

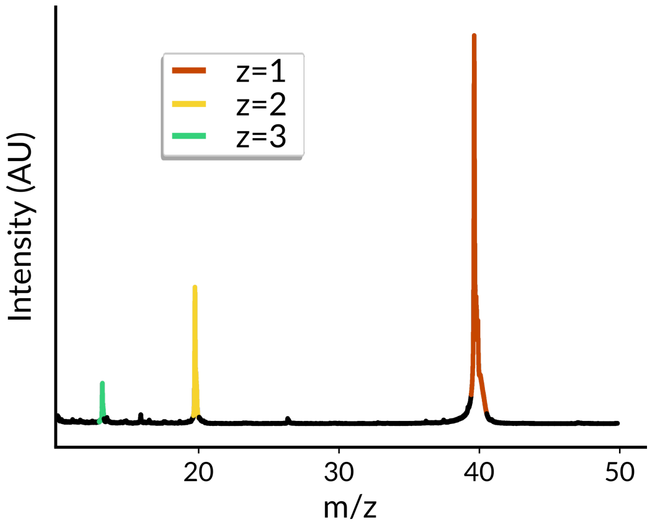


Figure S1: MALDI-TOF spectra of TR(1,11) protein. Peaks are labeled by color according to the appropriate mass-to-charge ratio.

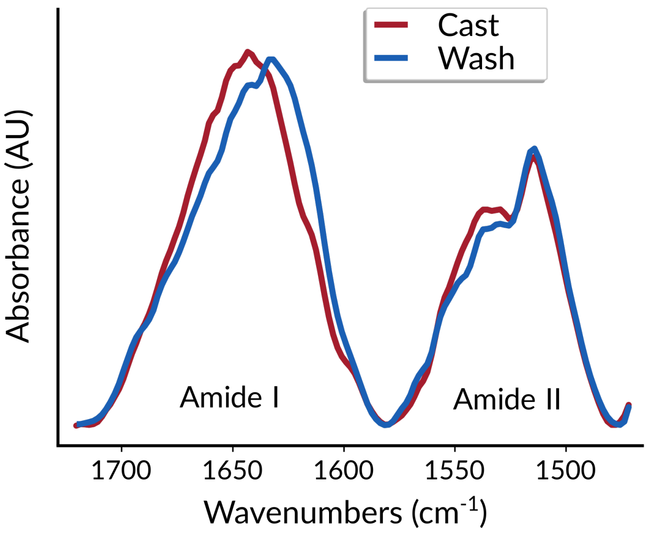


Figure S2: FTIR spectra of TR(1,11) amide I and II regions.

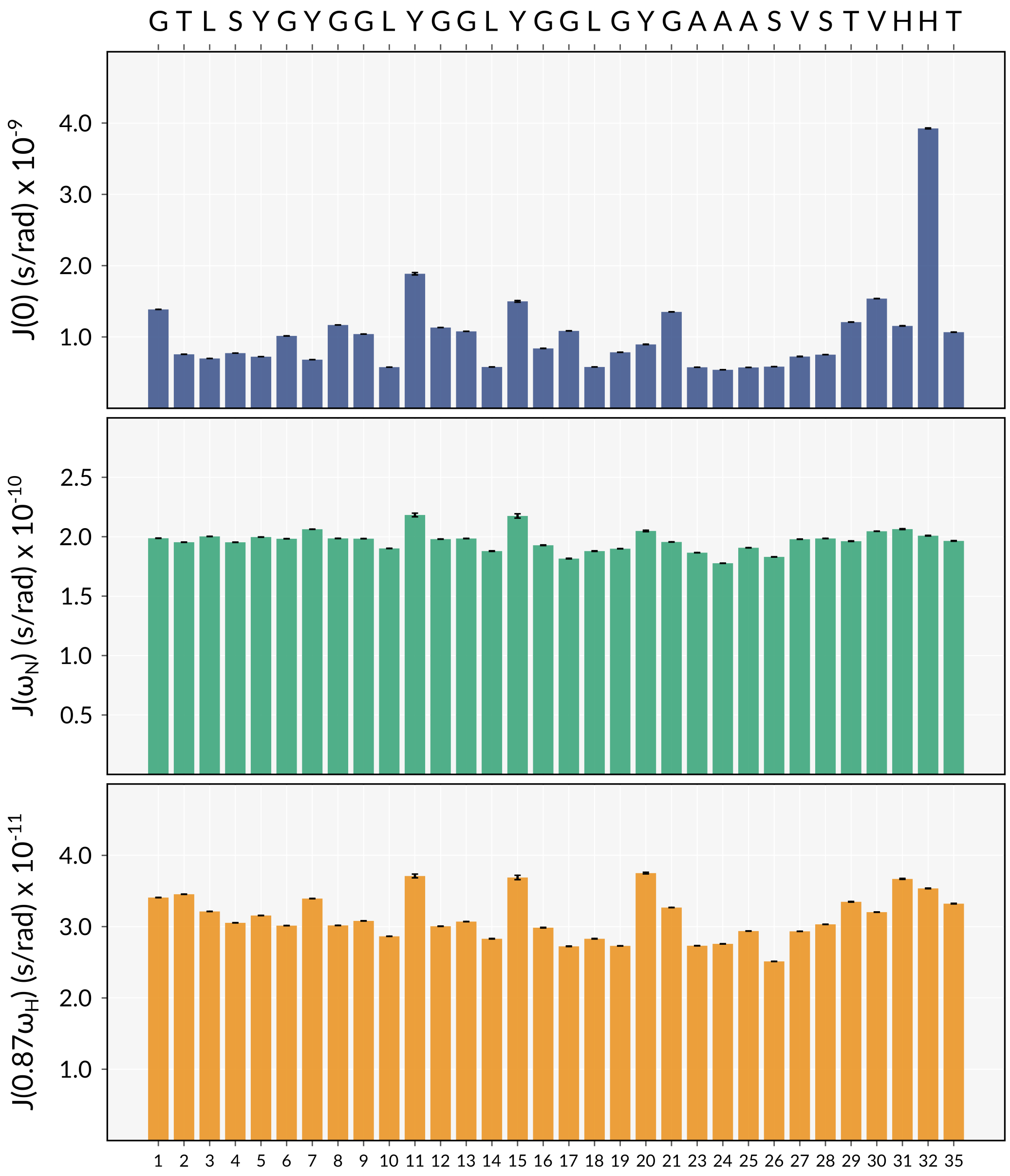


Figure S3: $J(0)$, $J(\omega_{\mathrm{N}})$ and $J(0.87\omega_{\mathrm{H}})$ spectral density values derived from the $^{15}$N relaxation rates and heteronuclear nOe experiments.

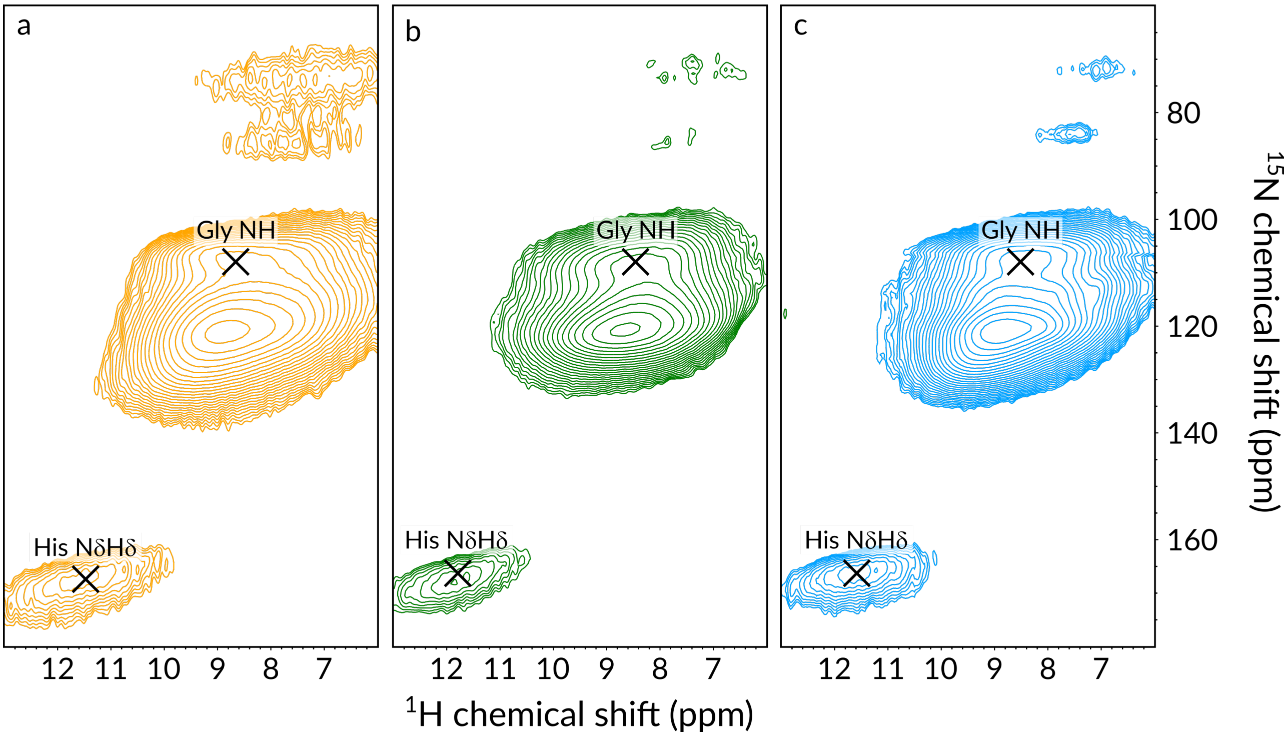


Figure S4: $^1$H-$^{15}$N CP-based 2D spectra of sample AP (a), AF (b) and HF (c) of $^{13}$C,$^{15}$N-labeled TR(1,11) recorded at 700 MHz $^1$H Larmor frequency at 55.55 kHz MAS.

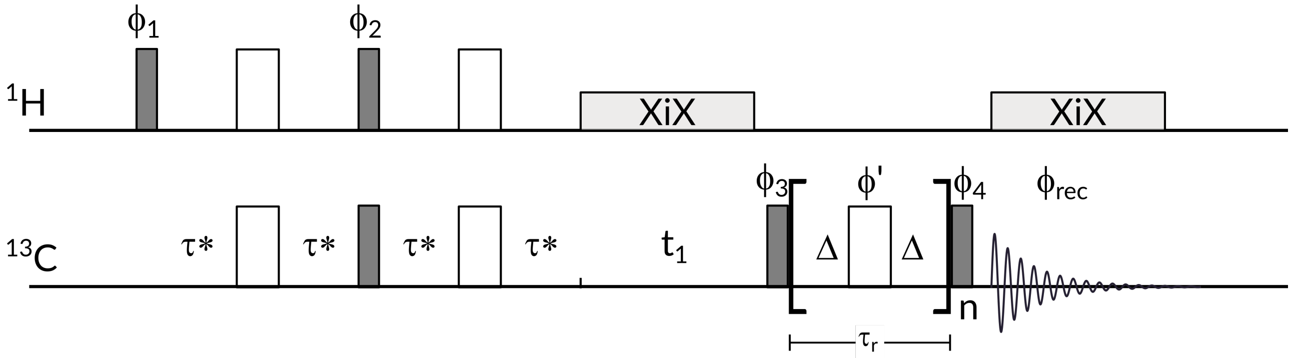


Figure S5: Pulse sequence of the refocused INEPT-based 2D $^{13}$C–$^{13}$C experiment using RFDR mixing for magnetization transfer. The initial $^{13}$C polarization is created via a refocused INEPT step where $\tau^* \approx 0.45(1/4J)$ in order to mitigate excessive coherence decay during the refocused INEPT step.[2] Following the indirect evolution period $t_1$, a non-selective $\pi/2$ generates longitudinal $^{13}$C magnetization and rotor-synchronized $\pi$ pulses are applied once per rotor period ($\tau_r$) for $n = 256$ times. $\pi$ pulses are phase-cycled according to the XY–16 scheme in order to minimize the pulse imperfections. An XiX decoupling is applied during the indirect and direct acquisition time. Pulse phases are $\phi_1 = x, -x$, $\phi_2 = 2(-y), 2(y)$, $\phi_3 = 4(y), 4(-y)$, $\phi_{\text{rec}} = x, -x, -x, x, -x, x, x, -x$; $\phi_4$ is decremented by 90° to achieve quadrature detection of $t_1$ according to the States-TPPI scheme.

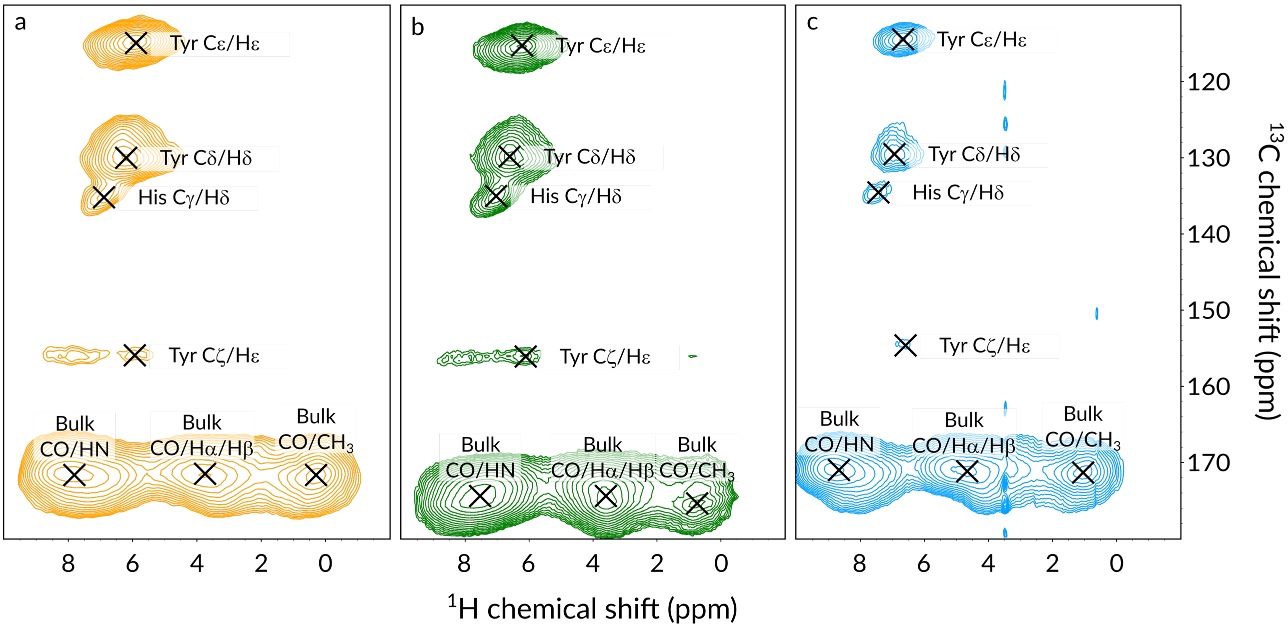


Figure S6: $^{1}$H-$^{13}$CO CP-based 2D spectra of sample AP (a), AF (b), and HF (c) of $^{13}$C,$^{15}$N-labeled TR(1,11) recorded at 700 MHz $^{1}$H Larmor frequency at 55.55 kHz MAS.

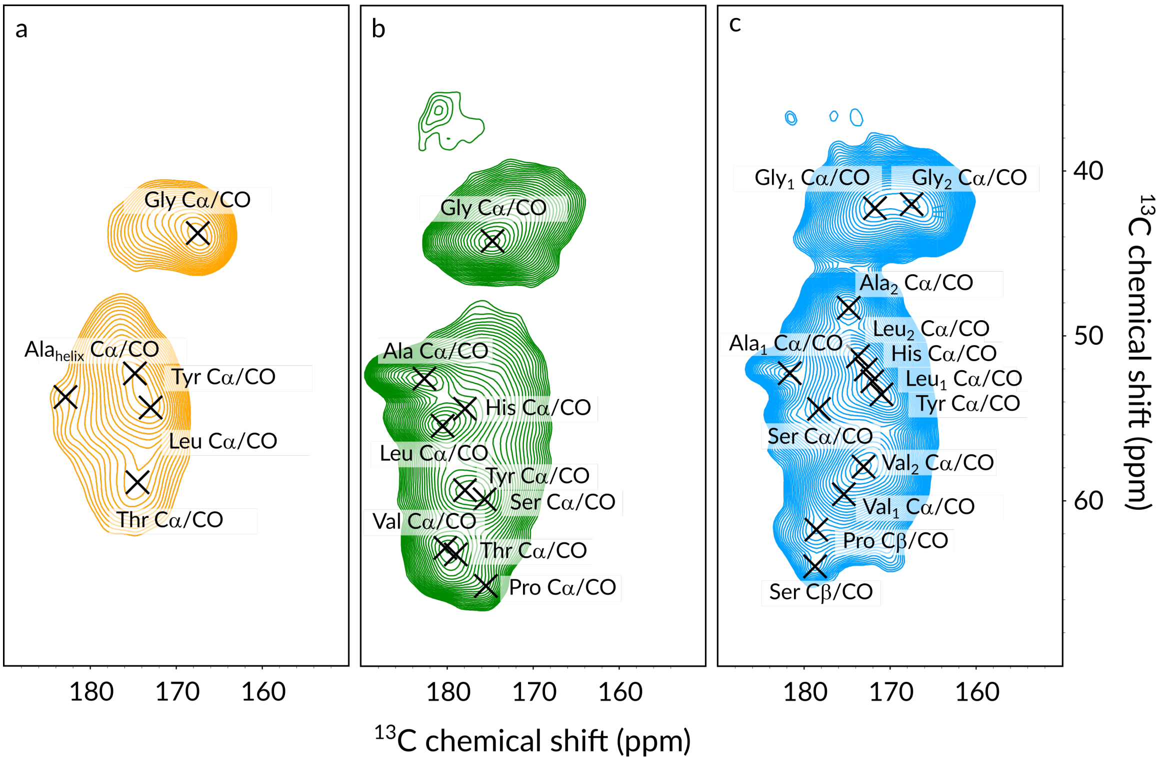


Figure S7: $^{1}$H-$^{13}$CO CP-based 2D spectra of sample AP (a), AF (b), and HF (c) of $^{13}$C,$^{15}$N-labeled TR(1,11) recorded at 700 MHz $^{1}$H Larmor frequency at 55.55 kHz MAS.

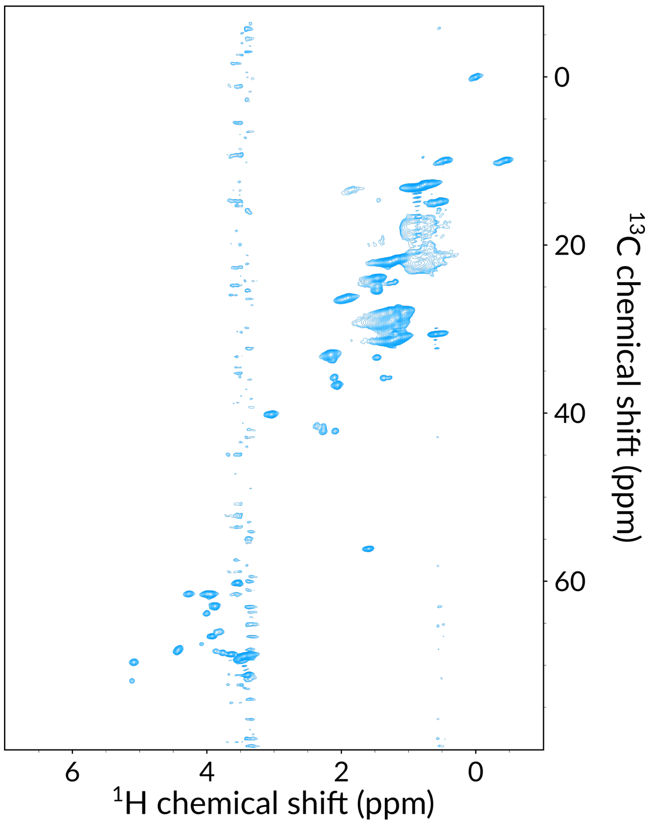


Figure S8: Excerpt from the aliphatic/methyl region of the $^{1}$H-$^{13}$C HSQC spectrum of sample HF of $^{13}$C,$^{15}$N-labeled TR(1,11) recorded at 700 MHz $^{1}$H Larmor frequency at 55.55 kHz MAS at ∼40°C.

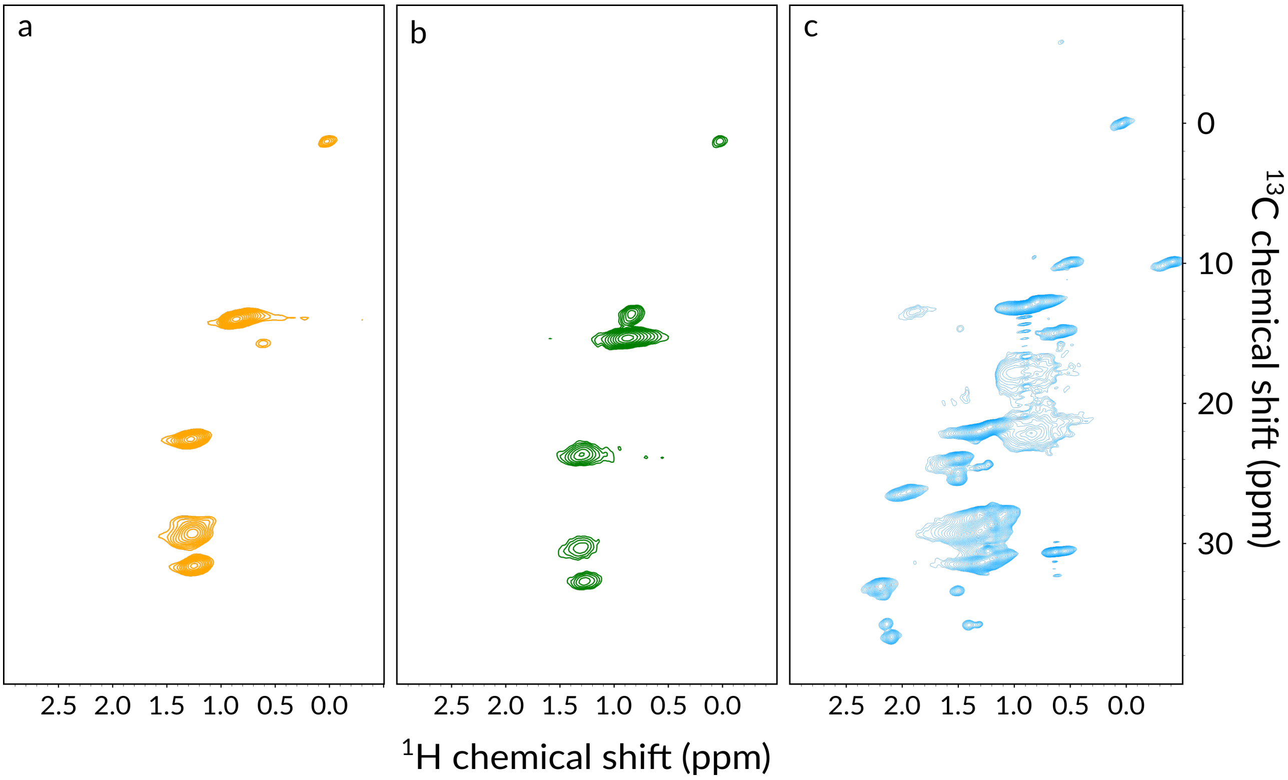


Figure S9: Comparison of the methyl region of the $^{1}$H-$^{13}$C HSQC spectrum of sample AP (a), AF (b), and HF (c) of $^{13}$C,$^{15}$N-labeled TR(1,11) recorded at 700 MHz $^{1}$H Larmor frequency at 55.55 kHz MAS.

Table S1: Deconvoluted FTIR spectral data for casted and washed films.

| **Assignment** | **Band position ($cm^{-1}$)** | **AF (%)** | **HF (%)** |
|---|---|---|---|
| $\alpha$-helix | 1658 – 1662 | 25.3 | 7.1 |
| $\beta$-sheet | 1630 – 1632/1697 – 1703 | 26.2 | 46.8 |
| Turn | 1665 – 1693 | 14.0 | 22.2 |
| Side–chain | 1594 – 1597 | 3.6 | 4.7 |
| Random Coil | 1640 – 1655 | 30.9 | 19.3 |

Table S2: $^{13}C$ chemical shift of TR(1,11) compared to literature values. The random coil, $\beta$-strand and $\alpha$-helix chemical shifts (labelled as "PACSY database") are reported as the statistical average for 3000+ folded proteins. Reference values are also reported for spider (*N. clavipes*) and silkworm (*B. mori*) silks ("Natural silks"). For all Pro entries, values for the *trans* isomer were reported. All shifts were directly referenced to DSS unless labelled with the * symbol, were conversion from TMS was achieved via addition of 1.7 ppm. Symbols "–" and "/" indicate a chemical shift range or multiple distinct environments, respectively.

| | | **PACSY database**[3] | | | **As spun silks** | | **Experimental TR(1,11) values** | | |
|---|---|---|---|---|---|---|---|---|---|
| **Residue** | **Index** | **Coil** | **$\beta$-strand** | **$\alpha$-helix** | **Dragline***[4,5] | ***B. Mori****[6] | **DMSO-$d_6$** | **AF** | **HF** |
| Gly $H^N$ | 1 | 8.34 | 8.35 | 8.28 | – | – | 8.06 | – | 8.4 |
| Gly $H_\alpha$ | | 3.93 | – | – | – | – | 3.72 – 3.82 | – | – |
| Gly CO | | 173.9 | 172.1 | 175.4 | 172.7 | 171.0 | 170.5 | 170.0 – 172.3 | 168.2/170.3 |
| Gly C$\alpha$ | | 45.4 | 45.1 | 46.9 | 45.0 | 44.2 | 43.4 | 42.2 – 45.0 | 41.6 |
| Gly $N^H$ | | 110.0 | 109.8 | 107.9 | – | – | 106.9 | – | 107.6 |
| Thr $H^N$ | 2 | 8.20 | 8.58 | 8.04 | – | – | 7.78/7.82 | – | – |
| Thr $H_\alpha$ | | 4.46 | – | – | – | – | 4.29 | – | – |
| Thr CO | | 174.5 | 173.6 | 176.0 | – | – | 171.2 | 174.4 | 171.1 |
| Thr C$\alpha$ | | 61.4 | 61.0 | 65.5 | – | – | 59.4 | 63.2 | 58.3 |
| Thr C$\beta$ | | 69.4 | 70.4 | 68.1 | – | – | 68.1 | 66.2 | 66.5 – 68.4 |
| Thr C$\gamma$ | | 21.6 | – | – | – | – | 20.8 | 18.0 | 18.3 |
| Thr $N^H$ | | 114.3 | 117.6 | 114.7 | – | – | 111.0 | – | – |
| Leu $H^N$ | 3 | 8.10 | 8.72 | 8.07 | – | – | 7.93 | – | – |
| Leu $H_\alpha$ | | 4.31 | – | – | – | – | 4.30 – 4.38 | – | – |
| Leu CO | | 176.4 | 175.4 | 178.2 | – | – | 173.5 | 175.5 | 170.7/171.5 |
| Leu C$\alpha$ | | 54.7 | 53.8 | 57.4 | – | – | 52.4 | 54.4 | 50.4/52.0 |
| Leu C$\beta$ | | 42.3 | 44.1 | 41.4 | – | – | 41.9 | 38.3 | 38.8/42.5 |
| Leu C$\gamma$ | | 26.9 | – | – | – | – | 25.3 | 23.1 | 23.3 – 23.7 |
| Leu C$\delta$ | | 24.5 | – | – | – | – | 24.3 | 18.4/21.4 | 21.5 |
| Leu $N^H$ | | 122.0 | 124.7 | 120.0 | – | – | 120.1 | – | – |
| Ser $H^N$ | 4 | 8.28 | 8.55 | 8.14 | – | – | 7.88 | – | – |
| Ser $H_\alpha$ | | 4.48 | – | – | – | – | 4.29/4.36 | – | – |
| Ser CO | | 174.4 | 173.5 | 176.0 | – | 173.9 | 171.4 | 173.1 | 173.9 |
| Ser C$\alpha$ | | 58.2 | 57.1 | 61.0 | 57.2 | 56.3 | 56.3 | 59.0 | 53.7 |
| Ser C$\beta$ | | 63.5 | 64.8 | 62.5 | 63.3 | 65.6 | 63.0 | 59.3 | 62.8 |
| Ser $N^H$ | | 116.3 | 117.4 | 115.1 | – | – | 113.3 | – | – |
| Tyr $H^N$ | 5 | 8.05 | 8.77 | 8.10 | – | – | 7.91 | – | – |
| Tyr $H_\alpha$ | | 4.61 | – | – | – | – | 4.45 – 4.51 | – | – |
| Tyr CO | | 174.9 | 174.3 | 177.1 | – | – | 173.0 | 173.9 | 170.1 |
| Tyr C$\alpha$ | | 57.4 | 56.6 | 60.6 | – | – | 55.8 | 58.4 | 53.0 |

| | | | | | | | | | |
|---|---|---|---|---|---|---|---|---|---|
| Tyr C$\beta$ | | 38.9 | 41.0 | 38.1 | – | – | 37.8 | 34.7 | 38.5 |
| Tyr C$\gamma$ | | 123.7 | – | – | – | – | – | 126.5 | 126.3 |
| Tyr C$\delta$ | 5 | 133.1 | – | – | – | – | 131.6 | 129.0 | 129.6 |
| Tyr C$\epsilon$ | | 117.9 | – | – | – | – | 116.3 | 114.7 | 114.4 |
| Tyr C$\zeta$ | | 157.6 | – | – | – | – | – | 154.7 | 154.6 |
| Tyr $N^H$ | | 118.9 | 121.6 | 119.6 | – | – | 118.4 | – | – |
| Gly $H^N$ | | 8.34 | 8.35 | 8.28 | – | – | 8.11 | – | 8.4 |
| Gly $H_\alpha$ | | 3.93 | – | – | – | – | 3.72 – 3.82 | – | – |
| Gly CO | 6 | 173.9 | 172.1 | 175.4 | 172.7 | 171.0 | 170.1 – 170.6 | 170.0 – 172.3 | 168.2/170.3 |
| Gly C$\alpha$ | | 45.4 | 45.1 | 46.9 | 45.0 | 44.2 | 43.4 | 42.2 – 45.0 | 41.6 |
| Gly $N^H$ | | 110.0 | 109.8 | 107.9 | – | – | 106.1 | – | 107.6 |
| Tyr $H^N$ | | 8.05 | 8.77 | 8.10 | – | – | 7.91 | – | – |
| Tyr $H_\alpha$ | | 4.61 | – | – | – | – | 4.45 – 4.51 | – | – |
| Tyr CO | | 174.9 | 174.3 | 177.1 | – | – | 173.0 | 173.9 | 170.1 |
| Tyr C$\alpha$ | | 57.4 | 56.6 | 60.6 | – | – | 55.8 | 58.4 | 53.0 |
| Tyr C$\beta$ | 7 | 38.9 | 41.0 | 38.1 | – | – | 37.8 | 34.7 | 38.5 |
| Tyr C$\gamma$ | | 123.7 | – | – | – | – | – | 126.5 | 126.3 |
| Tyr C$\delta$ | | 133.1 | – | – | – | – | 131.6 | 129.0 | 129.6 |
| Tyr C$\epsilon$ | | 117.9 | – | – | – | – | 116.3 | 114.7 | 114.4 |
| Tyr C$\zeta$ | | 157.6 | – | – | – | – | – | 154.7 | 154.6 |
| Tyr $N^H$ | | 118.9 | 121.6 | 119.6 | – | – | 118.4 | – | – |
| Gly $H^N$ | | 8.34 | 8.35 | 8.28 | – | – | 8.10 | – | 8.4 |
| Gly $H_\alpha$ | | 3.93 | – | – | – | – | 3.72 – 3.82 | – | – |
| Gly CO | 8 | 173.9 | 172.1 | 175.4 | 172.7 | 171.0 | 170.6 | 170.0 – 172.3 | 168.2/170.3 |
| Gly C$\alpha$ | | 45.4 | 45.1 | 46.9 | 45.0 | 44.2 | 43.1 | 42.2 – 45.0 | 41.6 |
| Gly $N^H$ | | 110.0 | 109.8 | 107.9 | – | – | 106.1 | – | 107.6 |
| Gly $H^N$ | | 8.34 | 8.35 | 8.28 | – | – | 8.12 | – | 8.4 |
| Gly $H_\alpha$ | | 3.93 | – | – | – | – | 3.72 – 3.82 | – | – |
| Gly CO | 9 | 173.9 | 172.1 | 175.4 | 172.7 | 171.0 | 170.2 | 170.0 – 172.3 | 168.2/170.3 |
| Gly C$\alpha$ | | 45.4 | 45.1 | 46.9 | 45.0 | 44.2 | 43.4 | 42.2 – 45.0 | 41.6 |
| Gly $N^H$ | | 110.0 | 109.8 | 107.9 | – | – | 105.7 | – | 107.6 |
| Leu $H^N$ | | 8.10 | 8.72 | 8.07 | – | – | 7.98 | – | – |
| Leu $H_\alpha$ | | 4.31 | – | – | – | – | 4.30 – 4.38 | – | – |
| Leu CO | | 176.4 | 175.4 | 178.2 | – | – | 173.4 | 175.5 | 170.7/171.5 |
| Leu C$\alpha$ | 10 | 54.7 | 53.8 | 57.4 | – | – | 52.5 | 54.4 | 50.4/52.0 |
| Leu C$\beta$ | | 42.3 | 44.1 | 41.4 | – | – | 42.1 | 38.3 | 38.8/42.5 |
| Leu C$\gamma$ | | 26.9 | – | – | – | – | 25.3 | 23.1 | 23.3 – 23.7 |
| Leu C$\delta$ | | 24.5 | – | – | – | – | 24.3 | 18.4/21.4 | 21.5 |
| Leu $N^H$ | | 122.0 | 124.7 | 120.0 | – | – | 118.0 | – | – |
| Tyr $H^N$ | | 8.05 | 8.77 | 8.10 | – | – | 7.95 | – | – |
| Tyr $H_\alpha$ | | 4.61 | – | – | – | – | 4.45 – 4.51 | – | – |
| Tyr CO | | 174.9 | 174.3 | 177.1 | – | – | 172.7 | 173.9 | 170.1 |
| Tyr C$\alpha$ | 11 | 57.4 | 56.6 | 60.6 | – | – | 55.7 | 58.4 | 53.0 |
| Tyr C$\beta$ | | 38.9 | 41.0 | 38.1 | – | – | 37.8 | 34.7 | 38.5 |
| Tyr C$\gamma$ | | 123.7 | – | – | – | – | – | 126.5 | 126.3 |
| Tyr C$\delta$ | | 133.1 | – | – | – | – | 131.6 | 129.0 | 129.6 |
| Tyr C$\epsilon$ | | 117.9 | – | – | – | – | 116.3 | 114.7 | 114.4 |

| | | | | | | | | | |
|---|---|---|---|---|---|---|---|---|---|
| Tyr Cζ | 11 | 157.6 | – | – | – | – | – | 154.7 | 154.6 |
| Tyr $N^H$ | | 118.9 | 121.6 | 119.6 | – | – | 116.6 | – | – |
| Gly $H^N$ | 12 | 8.34 | 8.35 | 8.28 | – | – | 8.11 | – | 8.4 |
| Gly $H_\alpha$ | | 3.93 | – | – | – | – | 3.72 – 3.82 | – | – |
| Gly CO | | 173.9 | 172.1 | 175.4 | 172.7 | 171.0 | 170.1 – 170.6 | 170.0 – 172.3 | 168.2/170.3 |
| Gly Cα | | 45.4 | 45.1 | 46.9 | 45.0 | 44.2 | 43.4 | 42.2 – 45.0 | 41.6 |
| Gly $N^H$ | | 110.0 | 109.8 | 107.9 | – | – | 106.1 | – | 107.6 |
| Gly $H^N$ | 13 | 8.34 | 8.35 | 8.28 | – | – | 8.12 | – | 8.4 |
| Gly $H_\alpha$ | | 3.93 | – | – | – | – | 3.72 – 3.82 | – | – |
| Gly CO | | 173.9 | 172.1 | 175.4 | 172.7 | 171.0 | 170.3 | 170.0 – 172.3 | 168.2/170.3 |
| Gly Cα | | 45.4 | 45.1 | 46.9 | 45.0 | 44.2 | 43.4 | 42.2 – 45.0 | 41.6 |
| Gly $N^H$ | | 110.0 | 109.8 | 107.9 | – | – | 105.7 | – | 107.6 |
| Leu $H^N$ | 14 | 8.10 | 8.72 | 8.07 | – | – | 7.95 | – | – |
| Leu $H_\alpha$ | | 4.31 | – | – | – | – | 4.30 – 4.38 | – | – |
| Leu CO | | 176.4 | 175.4 | 178.2 | – | – | 173.5 | 175.5 | 170.7/171.5 |
| Leu Cα | | 54.7 | 53.8 | 57.4 | – | – | 52.7 | 54.4 | 50.4/52.0 |
| Leu Cβ | | 42.3 | 44.1 | 41.4 | – | – | 41.7 | 38.3 | 38.8/42.5 |
| Leu Cγ | | 26.9 | – | – | – | – | 25.3 | 23.1 | 23.3 – 23.7 |
| Leu Cδ | | 24.5 | – | – | – | – | 24.3 | 18.4/21.4 | 21.5 |
| Leu $N^H$ | | 122.0 | 124.7 | 120.0 | – | – | 118.4 | – | – |
| Tyr $H^N$ | 15 | 8.05 | 8.77 | 8.10 | – | – | 7.95 | – | – |
| Tyr $H_\alpha$ | | 4.61 | – | – | – | – | 4.45 – 4.51 | – | – |
| Tyr CO | | 174.9 | 174.3 | 177.1 | – | – | 172.7 | 173.9 | 170.1 |
| Tyr Cα | | 57.4 | 56.6 | 60.6 | – | – | 55.7 | 58.4 | 53.0 |
| Tyr Cβ | | 38.9 | 41.0 | 38.1 | – | – | 37.8 | 34.7 | 38.5 |
| Tyr Cγ | | 123.7 | – | – | – | – | – | 126.5 | 126.3 |
| Tyr Cδ | | 133.1 | – | – | – | – | 131.6 | 129.0 | 129.6 |
| Tyr Cϵ | | 117.9 | – | – | – | – | 116.3 | 114.7 | 114.4 |
| Tyr Cζ | | 157.6 | – | – | – | – | – | 154.7 | 154.6 |
| Tyr $N^H$ | | 118.9 | 121.6 | 119.6 | – | – | 116.6 | – | – |
| Gly $H^N$ | 16 | 8.34 | 8.35 | 8.28 | – | – | 8.04 | – | 8.4 |
| Gly $H_\alpha$ | | 3.93 | – | – | – | – | 3.72 – 3.82 | – | – |
| Gly CO | | 173.9 | 172.1 | 175.4 | 172.7 | 171.0 | 170.1 – 170.6 | 170.0 – 172.3 | 168.2/170.3 |
| Gly Cα | | 45.4 | 45.1 | 46.9 | 45.0 | 44.2 | 43.4 | 42.2 – 45.0 | 41.6 |
| Gly $N^H$ | | 110.0 | 109.8 | 107.9 | – | – | 105.5 | – | 107.6 |
| Gly $H^N$ | 17 | 8.34 | 8.35 | 8.28 | – | – | 7.99 | – | 8.4 |
| Gly $H_\alpha$ | | 3.93 | – | – | – | – | 3.72 – 3.82 | – | – |
| Gly CO | | 173.9 | 172.1 | 175.4 | 172.7 | 171.0 | 170.3 | 170.0 – 172.3 | 168.2/170.3 |
| Gly Cα | | 45.4 | 45.1 | 46.9 | 45.0 | 44.2 | 43.4 | 42.2 – 45.0 | 41.6 |
| Gly $N^H$ | | 110.0 | 109.8 | 107.9 | – | – | 105.3 | – | 107.6 |
| Leu $H^N$ | 18 | 8.10 | 8.72 | 8.07 | – | – | 7.95 | – | – |
| Leu $H_\alpha$ | | 4.31 | – | – | – | – | 4.30 – 4.38 | – | – |
| Leu CO | | 176.4 | 175.4 | 178.2 | – | – | 173.8 | 175.5 | 170.7/171.5 |
| Leu Cα | | 54.7 | 53.8 | 57.4 | – | – | 52.7 | 54.4 | 50.4/52.0 |
| Leu Cβ | | 42.3 | 44.1 | 41.4 | – | – | 41.7 | 38.3 | 38.8/42.5 |
| Leu Cγ | | 26.9 | – | – | – | – | 25.3 | 23.1 | 23.3 – 23.7 |
| Leu Cδ | | 24.5 | – | – | – | – | 24.3 | 18.4/21.4 | 21.5 |

| | | | | | | | | | |
|---|---|---|---|---|---|---|---|---|---|
| Leu $N^H$ | 18 | 122.0 | 124.7 | 120.0 | – | – | 118.4 | – | – |
| Gly $H^N$ | 19 | 8.34 | 8.35 | 8.28 | – | – | 8.10 | – | 8.4 |
| Gly $H_\alpha$ | | 3.93 | – | – | – | – | 3.72 – 3.82 | – | – |
| Gly CO | | 173.9 | 172.1 | 175.4 | 172.7 | 171.0 | 170.1 | 170.0 – 172.3 | 168.2/170.3 |
| Gly C$\alpha$ | | 45.4 | 45.1 | 46.9 | 45.0 | 44.2 | 43.1 | 42.2 – 45.0 | 41.6 |
| Gly $N^H$ | | 110.0 | 109.8 | 107.9 | – | – | 105.6 | – | 107.6 |
| Tyr $H^N$ | 20 | 8.05 | 8.77 | 8.10 | – | – | 8.02 | – | – |
| Tyr $H_\alpha$ | | 4.61 | – | – | – | – | 4.45 – 4.51 | – | – |
| Tyr CO | | 174.9 | 174.3 | 177.1 | – | – | 173.1 | 173.9 | 170.1 |
| Tyr C$\alpha$ | | 57.4 | 56.6 | 60.6 | – | – | 55.8 | 58.4 | 53.0 |
| Tyr C$\beta$ | | 38.9 | 41.0 | 38.1 | – | – | 38.0 | 34.7 | 38.5 |
| Tyr C$\gamma$ | | 123.7 | – | – | – | – | – | 126.5 | 126.3 |
| Tyr C$\delta$ | | 133.1 | – | – | – | – | 131.6 | 129.0 | 129.6 |
| Tyr C$\epsilon$ | | 117.9 | – | – | – | – | 116.3 | 114.7 | 114.4 |
| Tyr C$\zeta$ | | 157.6 | – | – | – | – | – | 154.7 | 154.6 |
| Tyr $N^H$ | | 118.9 | 121.6 | 119.6 | – | – | 117.0 | – | – |
| Gly $H^N$ | 21 | 8.34 | 8.35 | 8.28 | – | – | 8.29 | – | 8.4 |
| Gly $H_\alpha$ | | 3.93 | – | – | – | – | 3.72 – 3.82 | – | – |
| Gly CO | | 173.9 | 172.1 | 175.4 | 172.7 | 171.0 | – | 170.0 – 172.3 | 168.2/170.3 |
| Gly C$\alpha$ | | 45.4 | 45.1 | 46.9 | 45.0 | 44.2 | 43.4 | 42.2 – 45.0 | 41.6 |
| Gly $N^H$ | | 110.0 | 109.8 | 107.9 | – | – | 106.6 | – | 107.6 |
| Pro $H_\alpha$ | 22 | 4.39 | – | – | – | – | – | – | – |
| Pro CO | | 176.5 | 176.0 | 178.0 | 176.5 | – | 172.8 | 173.1 | 173.9 |
| Pro C$\alpha$ | | 63.0 | 62.7 | 65.3 | 62.0 | – | 60.5 | 63.4 | 61.3 |
| Pro C$\beta$ | | 32.0 | 32.1 | 31.6 | 32.2 | – | 30.0 | 28.7 | 29.3 |
| Pro C$\gamma$ | | 27.3 | – | – | 27.1 | – | 25.1 | 23.8 | 23.7 |
| Pro C$\delta$ | | 50.3 | – | – | 49.2 | – | 47.0 | 46.1 | 46.3 |
| Ala $H^N$ | 23 | 8.19 | 8.60 | 8.10 | – | – | 8.10 | – | – |
| Ala $H_\alpha$ | | 4.25 | – | – | – | – | 4.24/4.34 | – | – |
| Ala CO | | 176.9 | 175.6 | 179.1 | 174.3 | 173.9 | 173.7 | 176.8 | 172.0/175.5 |
| Ala C$\alpha$ | | 52.4 | 50.9 | 54.7 | 49.9 | 50.6 | 49.5 | 51.7 | 47.8/51.8 |
| Ala C$\beta$ | | 19.4 | 21.7 | 18.5 | 22.6 | 18.3/21.3 | 19.3 | 14.0 | 14.4/19.5 |
| Ala $N^H$ | | 124.1 | 124.9 | 121.8 | – | – | 119.4 | – | – |
| Ala $H^N$ | 24 | 8.19 | 8.60 | 8.10 | – | – | 7.79 | – | – |
| Ala $H_\alpha$ | | 4.25 | – | – | – | – | 4.24/4.34 | – | – |
| Ala CO | | 176.9 | 175.6 | 179.1 | 174.3 | 173.9 | 173.5 | 176.8 | 172.0/175.5 |
| Ala C$\alpha$ | | 52.4 | 50.9 | 54.7 | 49.9 | 50.6 | 49.6 | 51.7 | 47.8/51.8 |
| Ala C$\beta$ | | 19.4 | 21.7 | 18.5 | 22.6 | 18.3/21.3 | 19.1 | 14.0 | 14.4/19.5 |
| Ala $N^H$ | | 124.1 | 124.9 | 121.8 | – | – | 118.3 | – | – |
| Ala $H^N$ | 25 | 8.19 | 8.60 | 8.10 | – | – | 7.93 | – | – |
| Ala $H_\alpha$ | | 4.25 | – | – | – | – | 4.24/4.34 | – | – |
| Ala CO | | 176.9 | 175.6 | 179.1 | 174.3 | 173.9 | 173.8 | 176.8 | 172.0/175.5 |
| Ala C$\alpha$ | | 52.4 | 50.9 | 54.7 | 49.9 | 50.6 | 49.5 | 51.7 | 47.8/51.8 |
| Ala C$\beta$ | | 19.4 | 21.7 | 18.5 | 22.6 | 18.3/21.3 | 19.3 | 14.0 | 14.4/19.5 |
| Ala $N^H$ | | 124.1 | 124.9 | 121.8 | – | – | 118.6 | – | – |
| Ser $H^N$ | 26 | 8.28 | 8.55 | 8.14 | – | – | 7.95 | – | – |
| Ser $H_\alpha$ | | 4.48 | – | – | – | – | 4.29/4.36 | – | – |

| | | | | | | | | | |
|---|---|---|---|---|---|---|---|---|---|
| Ser CO | 26 | 174.4 | 173.5 | 176.0 | – | 173.9 | 171.5 | 173.1 | 173.9 |
| Ser Cα | | 58.2 | 57.1 | 61.0 | 57.2 | 56.4 | 56.3 | 59.0 | 53.7 |
| Ser Cβ | | 63.5 | 64.8 | 62.5 | 63.3 | 65.6 | 62.9 | 59.3 | 62.8 |
| Ser $N^H$ | | 116.3 | 117.4 | 115.1 | – | – | 112.2 | – | – |
| Val $H^N$ | 27 | 8.06 | 8.70 | 8.00 | – | – | 7.63 | – | – |
| Val $H_\alpha$ | | 4.18 | – | – | – | – | 4.25 | – | – |
| Val CO | | 175.4 | 174.6 | 177.4 | – | – | 172.4 | 175.0 | 171.2/172.0 |
| Val Cα | | 61.7 | 60.7 | 65.8 | – | – | 58.7 | 61.8 | 57.5/59.1 |
| Val Cβ | | 32.8 | 33.9 | 31.5 | – | – | 31.9 | 27.9 | 28.9/31.5 |
| Val Cγ | | 21.5 | – | – | – | – | 20.5 | 17.5 | 17.3/17.7 |
| Val $N^H$ | | 120.5 | 122.6 | 119.5 | – | – | 114.6 | – | – |
| Ser $H^N$ | 28 | 8.28 | 8.55 | 8.14 | – | – | 8.09 | – | – |
| Ser $H_\alpha$ | | 4.48 | – | – | – | – | 4.29/4.36 | – | – |
| Ser CO | | 174.4 | 173.5 | 176.0 | – | 173.9 | 171.9 | 173.1 | 173.9 |
| Ser Cα | | 58.2 | 57.1 | 61.0 | 57.2 | 56.4 | 56.3 | 59.0 | 53.7 |
| Ser Cβ | | 63.5 | 64.8 | 62.5 | 63.3 | 65.6 | 62.9 | 59.3 | 62.8 |
| Ser $N^H$ | | 116.3 | 117.4 | 115.1 | – | – | 115.6 | – | – |
| Thr $H^N$ | 29 | 8.20 | 8.58 | 8.04 | – | – | 7.76/7.81 | – | – |
| Thr $H_\alpha$ | | 4.46 | – | – | – | – | 4.29 | – | – |
| Thr CO | | 174.5 | 173.6 | 176.0 | – | – | 171.6 | 174.4 | 171.1 |
| Thr Cα | | 61.4 | 61.0 | 65.5 | – | – | 59.6 | 63.2 | 58.3 |
| Thr Cβ | | 69.4 | 70.4 | 68.1 | – | – | 67.6 | 66.2 | 66.5 – 68.4 |
| Thr Cγ | | 21.6 | – | – | – | – | 20.8 | 18.0 | 18.3 |
| Thr $N^H$ | | 114.3 | 117.6 | 114.7 | – | – | 112.1 | – | – |
| Val $H^N$ | 30 | 8.06 | 8.70 | 8.00 | – | – | 7.65 | – | – |
| Val $H_\alpha$ | | 4.18 | – | – | – | – | 4.25 | – | – |
| Val CO | | 175.4 | 174.6 | 177.4 | – | – | 172.6 | 175.0 | 171.2/172.0 |
| Val Cα | | 61.7 | 60.7 | 65.8 | – | – | 59.3 | 61.8 | 57.5/59.1 |
| Val Cβ | | 32.8 | 33.9 | 31.5 | – | – | 31.3 | 27.9 | 28.9/31.5 |
| Val Cγ | | 21.5 | – | – | – | – | 20.5 | 17.5 | 17.3/17.7 |
| Val $N^H$ | | 120.5 | 122.6 | 119.5 | – | – | 115.6 | – | – |
| His $H^N$ | 31 | 8.22 | 8.70 | 8.00 | – | – | 7.95/8.27 | – | – |
| His $H_\alpha$ | | 4.60 | – | – | – | – | – | – | – |
| His CO | | 174.8 | 174.0 | 176.7 | – | – | 171.3 | 173.4 | 171.0 |
| His Cα | | 55.9 | 55.1 | 58.6 | – | – | 52.9 | 53.5 | 51.5 |
| His Cβ | | 30.2 | 32.2 | 30.0 | – | – | 28.0/30.2 | 27.4 | 29.3 |
| His Cδ | | 119.7 | – | – | – | – | 115.1 | 115.5 | 114.4 |
| His Cϵ | | 137.4 | – | – | – | – | 134.8 | – | – |
| His $N^H$ | | 119.7 | 121.9 | 118.0 | – | – | 118.8/119.7 | – | – |
| His $H^N$ | 32 | 8.22 | 8.70 | 8.00 | – | – | 8.10/8.31 | – | – |
| His $H_\alpha$ | | 4.60 | – | – | – | – | – | – | – |
| His CO | | 174.8 | 174.0 | 176.7 | – | – | – | 173.4 | 171.0 |
| His Cα | | 55.9 | 55.1 | 58.6 | – | – | 51.3/51.9 | 53.5 | 51.5 |
| His Cβ | | 30.2 | 32.2 | 30.0 | – | – | 27.3/29.2 | 27.4 | 29.3 |
| His Cδ | | 119.7 | – | – | – | – | 115.1 | 115.5 | 114.4 |
| His Cϵ | | 137.4 | – | – | – | – | 134.8 | – | – |
| His $N^H$ | | 119.7 | 121.9 | 118.0 | – | – | 117.9/119.7 | – | – |

| | | | | | | | | | |
|---|---|---|---|---|---|---|---|---|---|
| Pro $H_\alpha$ | 33 | 4.39 | – | – | – | – | – | – | – |
| Pro CO | | 176.5 | 176.0 | 178.0 | 176.5 | – | 173.7 | 173.1 | 173.9 |
| Pro $C\alpha$ | | 63.0 | 62.7 | 65.3 | 62.0 | – | 60.9 | 63.4 | 61.3 |
| Pro $C\beta$ | | 32.0 | 32.1 | 31.6 | 32.2 | – | 30.5 | 28.7 | 29.3 |
| Pro $C\gamma$ | | 27.3 | – | – | 27.1 | – | 25.8 | 23.8 | 23.7 |
| Pro $C\delta$ | | 50.3 | – | – | 49.2 | – | 47.5 | 46.1 | 46.3 |
| Ser $H^N$ | 34 | 8.28 | 8.55 | 8.14 | – | – | 8.38 | – | – |
| Ser $H_\alpha$ | | 4.48 | – | – | – | – | 4.29/4.36 | – | – |
| Ser CO | | 174.4 | 173.5 | 176.0 | – | 173.9 | 171.8 | 173.1 | 173.9 |
| Ser $C\alpha$ | | 58.2 | 57.1 | 61.0 | 57.2 | 56.4 | 56.8 | 59.0 | 53.7 |
| Ser $C\beta$ | | 63.5 | 64.8 | 62.5 | 63.3 | 65.6 | 62.7 | 59.3 | 62.8 |
| Ser $N^H$ | | 116.3 | 117.4 | 115.1 | – | – | 114.5 | – | – |
| Thr $H^N$ | 35 | 8.20 | 8.58 | 8.04 | – | – | 7.67 | – | – |
| Thr $H_\alpha$ | | 4.46 | – | – | – | – | 4.29 | – | – |
| Thr CO | | 174.5 | 173.6 | 176.0 | – | – | 171.9 | 174.4 | 171.1 |
| Thr $C\alpha$ | | 61.4 | 61.0 | 65.5 | – | – | 59.7 | 63.2 | 58.3 |
| Thr $C\beta$ | | 69.4 | 70.4 | 68.1 | – | – | 67.7 | 66.2 | 66.5 – 68.4 |
| Thr $C\gamma$ | | 21.6 | – | – | – | – | 20.8 | 18.0 | 18.3 |
| Thr $N^H$ | | 114.3 | 117.6 | 114.7 | – | – | 111.1 | – | – |

### References


1. Liu, J., Cao, D. & Zhang, L. Molecular dynamics study on nanoparticle diffusion in polymer melts: A test of the stokes-einstein law. *J. Phys. Chem. C* **112,** 6653–6661 (2008).
2. Rule, G. S. & Hitchens, T. K. *Fundamentals of protein NMR spectrosopy* (Springer, Dordrecht, The Netherlands, 2006).
3. Fritzsching, K. J., Hong, M. & Schmidt-Rohr, K. Conformationally selective multidimensional chemical shift ranges in proteins from a PACSY database purged using intrinsic quality criteria. *J. Biomol. NMR* **64,** 115–130 (2016).
4. Jenkins, J. E. *et al.* Solid-state NMR evidence for elastin-like $\beta$-turn structure in spider dragline silk. *Chem. Commun.* **46,** 6714–6716 (2010).
5. Jenkins, J. E., Holland, G. P. & Yarger, J. L. High resolution magic angle spinning NMR investigation of silk protein structure within major ampullate glands of orb weaving spiders. *Soft Matter* **8,** 1947–1954 (2012).
6. Asakura, T., Sugino, R., Yao, J., Takashima, H. & Kishore, R. Comparative structure analysis of tyrosine and valine residues in unprocessed silk fibroin (silk I) and in the processed silk fiber (silk II) from Bombyx mori using solid-state 13C,15N, and 2H NMR. *Biochemistry* **41,** 4415–4424 (2002).
